# *Limosilactobacillus fermentum* K73 synbiotic modulates gut microbiota function and behavior in gnotobiotic mice transplanted with microbiota from children diagnosed with autism spectrum disorder

**DOI:** 10.64898/2025.12.16.694732

**Authors:** Katherine Bauer Estrada, Karen M. Mancera Azamar, Caoyuanhui Wang, Samanvitha Deepthi Sudi, Kaleb Friday-Saunders, Victoria Márquez, Maria Ximena Quintanilla-Carvajal, Alejandro Acosta-González, Natalia Conde-Martinez, Ana Maria Porras, Karina Alviña

## Abstract

Dietary interventions targeting gut dysbiosis are gaining attention as potential therapeutic strategies for several pathological conditions. We previously demonstrated that an encapsulated synbiotic containing *Limosilactobacillus fermentum* K73 modulates human gut microbiota *in vitro*. Here, we extended this work using gnotobiotic mice colonized with gut microbiota from Colombian children diagnosed with autism spectrum disorder (ASD) or neurotypical (NT) donors. 16S rRNA sequencing showed that ASD- and NT-derived microbiota differed in both richness and composition, and humanized mice retained key features of their corresponding donor microbiota. Synbiotic supplementation significantly increased beneficial taxa abundance and short-chain fatty acid production, with more pronounced effects observed in ASD-colonized mice. These metabolic shifts were accompanied by behavioral improvements, including increased sociability and marble-burying performance, consistent with beneficial microbiota modulation. Together, these findings support synbiotic supplementation as a viable strategy to modulate dysbiotic gut microbiomes. Importantly, our results expand microbiome knowledge to understudied Latin American populations.

## 1. INTRODUCTION

Restoration of gut microbiota imbalance through dietary interventions has emerged as a potential therapeutic alternative for pathological conditions associated with gut dysbiosis (Larroya et al., 2021; Sonali et al., 2022; Cammann et al., 2023; Loh et al., 2024; Munoz-Pinto et al., 2024; Chatzinikolaou et al., 2025). Among these approaches, supplementation with synbiotics, which are mixtures of probiotics with fermentable prebiotic compounds, seem particularly promising to effectively modulate the gut microbiota due to the synergistic effects between their components that can enhance their overall impact (Swanson et al., 2020). Recent clinical studies have supported the therapeutic potential of synbiotics in pediatric populations diagnosed with ASD. For example, treatment with a synbiotic formulation comprised of guar gum and probiotics from the *Lactobacillus* and *Bifidobacterium* genera successfully modulated the gut microbiota of children diagnosed with autism spectrum disorder (ASD) (Mitchell et al., 2026). Similarly, synbiotics made with a mixture of probiotics and compounds such as epigallocatechin gallate and cranberry powder, among others, led to increased abundance of beneficial gut taxa, and improvements in gastrointestinal symptoms, language deficits, and speech patterns in ASD patients (Phan et al., 2024).

Building on this clinical evidence, we conducted an *in vivo* pilot study to test the potential modulatory effect of an encapsulated synbiotic comprised of high-oleic palm oil and *Limosilactobacillus fermentum* (*L. fermentum*) K73, isolated from the artisanal production of “Costeño” Cheese in Colombia (Cueto & Aragón, 2012; Bauer Estrada, Cardoso Cardenas, et al., 2025). This probiotic has a robust hypocholesterolemic effect due to the presence and activity of a bile salt hydrolase enzyme (Cardoso-Cardenas et al., 2025; Cueto & Aragón, 2012). Furthermore, we have recently shown that this encapsulated synbiotic can directly modulate human-derived gut microbiota in an *in vitro* batch bioreactor leading to increased production of short-chain fatty acids (SCFAs) and relative abundance of beneficial microorganisms (Bauer Estrada, Conde-Martínez, et al., 2025). These results support the *L. fermentum* K73 synbiotic formulation as a promising candidate for further *in vivo* evaluation of its potential therapeutic effect to modulate the gut microbiome.

For this pilot study, we focused on testing the impact of the *L.* fermentum K73 synbiotic on gut microbiota derived from children diagnosed with ASD for several reasons. First, in addition to having social interaction challenges, ASD patients often exhibit hyper- or hypo-reactivity to sensory stimuli (American Psychiatric Association, 2013) that can influence feeding behaviors, frequently resulting in pronounced food selectivity (Marí-Bauset et al., 2014). This can negatively affect their overall diet quality and nutritional status, factors contributing to altered gut microbiota composition (Putignani et al., 2014). Second, recent studies have shown that children with ASD have an imbalanced gut microbiota compared to neurotypical (NT) peers (Finegold et al., 2010; Hsiao et al., 2013; Kang et al., 2013; Ho et al., 2020; Bojović et al., 2020). Third, gut dysbiosis and modified dietary patterns result in distinct metabolic profiles characterized by altered SCFA and neurotransmitter production (Plaza-Díaz et al., 2019; Taniya et al., 2022; Morton et al., 2023). Hence, we sought to assess the efficacy of our *L. fermentum* K73 synbiotic for microbiome modulation using a gnotobiotic mouse model colonized with gut microbiota from Colombian children with or without ASD diagnosis. We selected a gnotobiotic mouse model because it allows for the controlled evaluation of microbiome-targeted interventions by enabling colonization with defined microbial communities (Grover & Kashyap, 2014; Luczynski et al., 2016). In fact, previous studies have shown that germ-free mice colonized with gut microbiota from children with ASD exhibit behavioral phenotypes that differ from those of mice colonized with microbiota from NT children (Sharon et al., 2019; Tengeler et al., 2020; Xiao et al., 2021; Avolio et al., 2022).

Here, we show that treatment with the *L. fermentum* synbiotic resulted in significant changes in gut microbiome function in gnotobiotic male mice colonized with ASD-associated gut microbiota compared to mice colonized with microbiota from NT donors. Specifically, the synbiotic treatment increased beneficial SCFAs and serotonin production only in mice transplanted with ASD-associated microbiota, with concomitant behavioral changes that mirrored these beneficial gut microbiome functional changes. Our results also indicate that humanized germ-free mice are a valuable preclinical platform for assessing the safety and translational potential of synbiotics for future human applications. Additionally, they provide evidence of the successful colonization of gnotobiotic mice using gut microbiota from Colombian children, a population understudied in both microbiome and ASD research.

## 2. METHODS

### 2.1.1 Fecal donor recruitment process and inclusion/exclusion criteria

Collection of human fecal samples for this study was approved by The Ethical Committee of the Universidad de La Sabana and Clinica Universidad de La Sabana (Act of session No. 100). The research activities of this study were performed following the Declarations of Helsinki for the Ethical Principles for Medical Research Involving Human Participants. Informed consent was obtained from the parents or legal guardians of all participants.

Fecal samples were obtained from Colombian children previously diagnosed with ASD (N=7) by their treating physicians prior to enrollment in our study. ASD diagnosis must be made according to the Colombian Ministry of Health’s Clinical Protocol for ASD diagnosis and treatment, which explicitly recommends to use of the criteria provided by the DSM-5 (Diagnostic and Statistical Manual of Mental Disorders, Fifth Edition), the official handbook published by the American Psychiatric Association. These patients were recruited in collaboration with the Apushi Foundation for Childhood Development, a specialized health care and educational center for children with neurodevelopmental disorders located in Bogotá, Colombia. Importantly, while the Apushi Foundation does not diagnose patients with ASD or any other neurodevelopmental condition, they are dedicated to the care and education of patients with significant behavioral needs. During their time at Apushi, all children received the same standard diet typical of the region, with no reported dietary restriction or special regimens. Inclusion/exclusion criteria for participation in our study included an age range of 5-10 years old, considering that ASD diagnosis is reliable and unlikely to change after 5 years of age (Zwaigenbaum et al., 2009), and gut microbiota at this age resembles adult-like composition (Schoultz et al., 2025). We also established as requirement for participation the absence of gastrointestinal symptoms (including diarrhea) and antibiotic consumption for at least 6 months before sample collection. Given the vulnerability of this pediatric population, and in accordance with our approved protocols, clinical history beyond diagnosis, comorbidities, and the stated inclusion/exclusion criteria was not collected.

Comparable fecal samples were collected from age and gender-matched children that did not have ASD or any other neurodevelopmental disorder diagnosis following the same inclusion/exclusion criteria as ASD donors. These samples are referred to as neurotypical or NT throughout our study (N=18). NT donors were recruited by publicly advertising our study at a local public school; therefore, while they lived in the same region as ASD donors, NT donors were not associated with the Apushi Foundation. Moreover, we included a larger number of NT samples to appropriately reflect the higher variability expected in gut microbiome composition among NT donors, who were recruited from a broader range of home environments and daily routines compared to the more uniform setting shared by ASD donors at the Apushi Foundation.

### 2.1.2. Human fecal sample collection and processing

Collection kits were provided to the parents/legal guardians with clear instructions on how to obtain clean fecal samples. Sample collection was carried out the night or morning before the samples were delivered to the care center and kept refrigerated at 4 °C for preservation. Upon arrival at the laboratory, samples were stored using the fecal aliquot straw technique (FAST) method as previously described(Romano et al., 2018) (136) and immediately stored at −80 °C. DNA was extracted from 100 mg of fecal matter using the QIAamp PowerFecal Pro DNA kit (Qiagen, Germany) following the kit instructions. The quantity of DNA was assessed using Qubit and A260/230 and 260/280 ratios were measured on a NanoDrop®1000 (Thermo Fisher Scientific, Waltham, MA, USA). Extracted DNA was stored at −20 °C for further analysis.

### 2.2 Colonization of gnotobiotic mice with human-associated gut microbiota

Fecal samples from Colombian donors were shipped frozen at −80 °C to the University of Florida (IRB protocol #IRB202302069). This study is covered under the ANLA authorization #3609 and the agreement granted by the Colombian Ministry of Environment and Sustainable Development “Contract for Access to Genetic Resources and their derivative products” (“*Contrato Marco de Acceso a Recursos Genéticos y sus Productos Derivados*”, #324 −2021 RGE0381).

For colonization, germ-free (GF) male mice (C57BL/6 wildtype strain, n=32) were bred in the GF facility at the University of Florida (IACUC protocol #202100000052). At 4-6 weeks old, these mice were securely transferred to a BSL-2 facility for subsequent inoculation and housing throughout the experiment, during which they were kept in sterile ventilated cages with autoclaved bedding material, and with *ad libitum* access to water and standard chow (2018x, Inotiv, USA). The room temperature was kept constant at 24 °C and 35% humidity, and the lights were kept on a 12/12 cycle (lights ON at 7AM, OFF at 7PM). Eight donor samples were used to humanize GF mice: four from donors with ASD diagnosis and four without (referred to as neurotypical or NT). 3-5 mice per donor sample were inoculated via gavage immediately upon arrival, using a 200 µL sample of fecal slurry prepared in phosphate buffered saline (PBS) under anaerobic conditions(Porras et al., 2021). Mice inoculated with the same donor slurry were kept in the same cage throughout the study. Fecal samples were collected at the start of the experiment and then weekly after inoculation for assessment of mouse gut microbiome composition. Food consumption and body weight were monitored every other day throughout the study.

### 2.3 Preparation of *L. fermentum* K73 synbiotic

Powdered synbiotic containing *L. fermentum* K73, its fermentation metabolites, high oleic palm oil and whey was prepared as we have previously described (Bauer Estrada, Conde-Martínez, et al., 2025). Briefly, *L. fermented* K73 was grown in food grade media and concentrated by centrifugation and then mixed with high oleic palm oil and lecithin for the first water-in-oil (W/O) phase. Separately, whey was homogenized with water and metabolites resulted from probiotic fermentation, using an Ultraturrax (IKA, US) and then mixed slowly with the W/O phase to form water-in-oil-in-water (W/O/W). Then, W/O/W emulsion was sealed by spray drying obtaining finally the synbiotic in powder. The synbiotic carrier was prepared following the same procedure without mixing *L. fermentum* K73 in.

### 2.4 Optimization of synbiotic administration in wildtype mice

All animal procedures described were previously approved by the University of Florida IACUC office (protocol #202100000052). Adult (4-6 months old) wildtype male mice (C57BL/6 background strain) were divided in groups of 5 per cage and received the following dietary treatments: (1) *ad libitum* standard chow (Std) – the control group, (2) carrier-only of bread-like food (CarBread), (3) synbiotic incorporated into the same bread-like food as Group 2 (SynBread), (4) carrier-only in powdered form (CarPowder), and (5) synbiotic in powdered form (SynPowder). Bread-like food was prepared by mixing 79.2% of wheat Flour, 0.5% of baking powder, 2.8% of margarine, 8% of water and 10% of powdered synbiotic or carrier. This mixed dough was then baked at 180 °C for 15 minutes and cut it into small pieces, resembling standard chow pellets.

Dietary synbiotic treatments were administered for 21 consecutive days before behavioral testing, during which overall health was monitored daily, and body weight was measured every other day. Mice fed with bread-like food containing either carrier only or carrier and synbiotic received 10 g of food every other day (10% was bread-like food and 90% was regular standard chow). Similarly, mice receiving powdered synbiotic were given 10 g of food in total, from which 1g (10% w/w) corresponded to synbiotic (7.2 logCFU/g) or carrier thoroughly mixed with regular standard chow pellets. The nutritional value of the synbiotic and its carrier was calculated as follows: total carbohydrates 63.2%, total fat 11.5%, total protein 12.2%, 10% probiotic (for synbiotic) and water 3.1%. With this the caloric intake was 13.8 kcal for synbiotic and carrier in powder treatments and 16.4 kcal for synbiotic and carrier in bread treatments. For both groups (synbiotic and carrier only), food was replenished every other day, and total consumption was measured by weighing any remaining food before replacing it with a fresh batch.

### 2.5 Treatment of colonized mice with *L. fermentum* K73 synbiotic

Treatment with *L. fermentum* K73 synbiotic was initiated after 4 weeks of inoculation with human-associated microbiota and one week of pre-synbiotic behavioral testing. The synbiotic was then administered in powdered form as described for WT mice (10% w/w in standard chow) for 21 days and continuously during post-synbiotic behavioral testing (28 days in total). The synbiotic-containing food was replaced every other day to ensure aseptic conditions and to carefully monitor the total amount consumed by weighing any remaining food before replacement.

### 2.6 Mouse fecal sample collection and processing

Mice were placed in a sterile plastic beaker each week for weighing and fecal sample collection. The fecal samples were placed in sterile microcentrifuge tubes, flash frozen, and stored in the laboratory at −80 °C. DNA was extracted from fecal samples collected at 4 timepoints: (1) at the start of the experiment upon GF mice arrival, (2) 4 weeks after inoculation, (3) at the start of the synbiotic treatment and (4) 3 weeks after synbiotic administration. DNA was extracted using the DNEasy Soil kit (Qiagen, Germany) and its concentration and A260/230 ratio were measured using Nanodrop (ThermoFisher, USA). DNA was stored at −20 °C for further analysis.

### 2.7 Gut microbiome analysis

The V3-V4 region of the 16S rRNA gene was amplified using the specific primers CCTAYGGGRBGCASCAG and GGACTACNNGGGTATCTAAT (Novogene, USA). PCR products were quantified and qualified by electrophoresis on a 2% agarose gel, and the mixture of PCR products was purified with a Qiagen Gel Extraction Kit (Qiagen, Germany). Sequencing libraries were generated using TruSeq® DNA PCR-Free Sample Preparation Kit (Illumina, USA) following the kit instructions and Novogene protocols for sequencing on an Illumina NovaSeq platform (Novogen, USA).

Paired-end reads were assigned to each sample by its unique barcode. Then, this barcode was removed and reads trimmed by cutting off the primers used. The paired end reads per sample were merged using FLASH V1.2.11 (Magoč & Salzberg, 2011), and quality filtering was performed using fastp 0.23.1 software to obtain Clean Tags(Bokulich et al., 2013). The tags were compared with the Silva reference database (https://www.arb-silva.de/) using the UCHIME Algorithm (http://www.drive5.com/usearch/manual/uchime_algo.html) to detect and remove chimera sequences(Edgar et al., 2011), obtaining the effective tags. Quality control of effective tags was carried out using FastQC to determine where reads should be truncated. The QIIME2 pipeline was followed starting with DADA2 denoising, where the reads were truncated to 240 bp. Then, taxonomy assignments were performed at 99% similarity using the SILVA database classifier for stool samples for the V3-V4 region(Quast et al., 2013). The phylogenetic tree was then calculated for this data set. Data frames of amplicon sequence variants (ASV) frequency per sample, sample metadata, and taxa assignment per ASV were obtained and analyzed using R software (V 2023.12.1). Later, a search of significantly different ASVs between group samples was carried out using the NCBI Nucleotide database in which the sequences of each ASV were compared to those of the 16S ribosomal RNA sequences, identifying the closest cultivable microorganism and the closest relative metagenome with a query of 100%. The frequency table, the taxonomy of the ASVs, and the phylogenetic tree were used to carry out the subsequent statistical analysis on R.

For all microbiota analyses, alpha diversity was calculated using the phyloseq richness function. Statistical analysis was performed according to data normality, using ANOVA or Kruskal-Wallis test as applicable using Benjamini-Hochberg correction for multiple testing. Beta diversity was calculated using Bray-Curtis distance matrices, followed by permutational multivariate analysis of variance (PERMANOVA) for comparison of different groups and corroborated with the beta-dispersity test. For the human donor samples, 2,532 different sequences were obtained, which corresponded to 3,252,000 reads, with a minimum of 29,100 and a maximum of 280,000 reads per sample. These readings were organized by ASV and per sample, obtaining a microbial composition frequency table where 2,525 ASVs were identified as bacteria with different lineages. ASVs identified as not belonging to the Bacteria domain were excluded. As not all samples had the same depth, they were rarefied to 28,000 reads per sample before assessing microbial diversity. Differential analysis was carried out using DESeq2 package. For both WT and GF mice, 2,736 different ASVs were obtained that corresponded to 9,050,249 reads with a minimum of reads per samples of 88,446. The samples were rarefied to a depth of 80,000 reads to perform all the analysis. Differential analysis was carried out using MaAsLin2 (Mallick et al., 2021) in R using a linear model followed by a t-test with a 95% confidence interval.

### 2.8 PCR analysis of the *L. fermentum K73 bsh* gene

To confirm that the *L. fermentum K73* probiotic was present in murine fecal samples, we performed end-point PCR for the *bsh* gene on selected fecal DNA samples collected before and after treatment with synbiotic. The *bsh* gene present in the *L.* fermentum genome encodes for the bile salt hydrolase enzyme capable of deconjugating bile acids. End-point PCR was performed using the primers LfHSB-f 5’-CCCAAGCCGCCTACCCCTCT-3’ Tm=64.3 °C and LfHSB-r 5’-CGGCAGGTGCGCCATCTTCA-3’ Tm=63.2 °C. The PCR was performed in a thermocycler (Eppendorf, USA) following standard protocols. PCR products were quantified using QuibitX (Thermo Fisher, USA) and observed in a 3% agarose gel.

### 2.9 Short chain fatty acid (SCFA) quantification

The production of acetic, butyric and propionic SCFAs was quantified as previously described (Bauer Estrada, Conde-Martínez, et al., 2025). Briefly, fecal samples were weighed and solubilized in 550 µL 0.1M phosphate buffer (pH = 7), reduced to pH = 3 (using 6M HCl solution) and then centrifuged at 10,000 g for 25 minutes at 4 °C. The supernatant was then filtered through a 0.22 μm nylon filter and loaded into an HPLC-DAD 3000 Ultimate (Thermo Fisher, USA), used with a SynergiTM 4 µm Hydro-RP 80Å column (250 x 4.6 mm) and a mobile phase starting at 100% water with H3PO4 at 0.1%, changing in a slow gradient to 20% acetonitrile. SCFAs were observed at 210 nm and integration of the area under the curve of each peak was performed for quantification using standards of each SCFA (Sigma-Aldrich, USA).

### 2.10 Quantification of serotonin levels

Serotonin levels were measured in mouse serum and fecal samples. Whole blood was collected from each mouse at the time of euthanasia, centrifuged at 2,000 g for 10 minutes at 4 °C, and the resulting serum was stored at −20 °C until analysis. Fecal pellets were weighed, solubilized in 550 µL phosphate buffer (pH = 7) and centrifuged at 10,000 g for 25 minutes at 4 °C to remove debris. The supernatant was collected and stored at −80 °C until analysis. Serotonin concentration was then quantified in both the serum and solubilized fecal samples using a commercial ELISA Kit (Enzo, USA), following the manufacturer’s instructions. Fecal and serum supernatant samples were diluted 1:2 and 1:90, respectively, in assay buffer prior to analysis. A standard curve was generated from serial dilutions (0.49 to 500 ng/mL) of the serotonin standard provided by the kit, and absorbance was read at 405 nm on a SpectraMax iD3 plate reader (Molecular Devices, USA). For fecal samples results were expressed as nM serotonin, normalized to fecal sample weight (nM/mg). For serum levels, results were expressed as nM concentration.

### 2.11 Colon Histology

After all behavioral testing was conducted, mice were deeply anesthetized with isoflurane and quickly euthanized prior to dissecting the colon. Any remaining stool was removed, and a 1 cm long section of the terminal colon was placed in a microcentrifuge tube containing 10% formalin. Fixed colon tissue samples were then sent to the Molecular Pathology Core at the University of Florida for subsequent paraffin embedding, sectioning, and staining with Hematoxylin & Eosin (H&E), following standard histological procedures. Colorimetric images of H&E-stained colon sections were obtained on a Keyence BZ-X800 microscope.

### 2.12 Behavioral analysis

Social, repetitive and exploratory behaviors were monitored before and after synbiotic treatment by conducting 3 different well-established tests: open field, marble burying and three-chamber test. To avoid excessive handling and potential contamination, only one behavioral test was performed per day, in the same BSL-2 room where mice were housed. All tests were performed at the same time each day to avoid circadian effects on behavior.

<u>Open field test (OFT):</u> The OFT was conducted in a square-shaped white plastic arena measuring 45.72 cm x 45.72 cm. Individual mice were placed in the center of the arena and allowed to explore for 10 minutes while video recording with a top-mounted camera. After each mouse, the arena was cleaned and disinfected with Peroxygard^TM^ (Virox Technologies, USA). EthoVision XT15 software (Noldus, The Netherlands) was used for video tracking analysis. The OFT arena was divided into two zones: the periphery and the central area (40% of the total area). For analysis, total distance, total time spent in each zone and the frequency of entries in each zone were quantified (Seibenhener & Wooten, 2015a).

<u>Marble burying test (MBT):</u> The MBT was performed using a clean autoclaved mouse cage (one per mouse), filled with autoclaved bedding material and levelled to 3 cm height. For the test, a total of 21 clean marbles (made of ceramics or wood) were placed on the surface of the bedding layer, arranged in an identical grid pattern of 7 rows of 3 marbles for each individual test. Prior to the start of each test, a photo was taken depicting all neatly organized marbles, then an individual mouse was placed in the center and allowed undisturbed exploration for 30 minutes. At the end of this period, each mouse was carefully removed and returned to their home cage, and a new photo was taken to document the placement of marbles left in the cage. The number of buried marbles, defined as >50% of total area covered by bedding, was quantified using ImageJ software (Angoa-Pérez et al., 2013). After each experiment, all marbles were collected and cleaned with Peroxygard^TM^ (Virox Technologies, USA) to disinfect and remove any remaining odor.

<u>Three Chamber Test (TCT):</u> The TCT was performed in a large white plastic rectangular arena measuring 44.5 cm x 59.7 cm. This arena was divided into three equal size chambers using dividers that had a small opening in the bottom to allow mice to move freely between chambers. In each of the lateral chambers a small container made of metal mesh was placed to hold either a novel object (nonsocial chamber) or a cage mate mouse (social chamber), while the center chamber was left empty. For the test, each mouse was placed in the center chamber and allowed to explore for 10 minutes while video recording. Videos were then analyzed using Ethovision XT15 software (Noldus, The Netherlands), and the time spent interacting with either the novel object or cage mate was quantified(Rein et al., 2020). We then calculated a sociability index (SI) according to the following equation: SI=Time in social chamber/ (Time in nonsocial chamber + Time in social chamber + Time in center). After each use, the arena was thoroughly cleaned using Peroxygard^TM^ (Virox Technologies, USA).

### 2.13 Statistical Analysis

Statistical analysis for SCFAs and serotonin measurements, and behavioral tests were conducted in R. ANOVA and Kruskal-Wallis tests with 95% confidence intervals were used to analyze normal and non-normally distributed data reported in this study. Dunn’s test for multiple comparisons was also performed to compare between all groups, and FDR corrections were performed for multiple groups testing. Finally, correlation analyses were performed using Spearman correlation.

## 3. RESULTS

### 3.1 Characterization of the gut microbiome of Colombian children with and without ASD diagnosis

First, we obtained fecal samples from children previously diagnosed with ASD, and age- and gender-matched neurotypical (NT) donors who had not been diagnosed with ASD or any other neurodevelopmental condition. Microbial analysis via 16S rRNA sequencing revealed that alpha diversity indices like observed features and Chao1 were significantly lower in children diagnosed with ASD compared to NT (p=0.038 and 0.03, respectively) while Shannon and phylogenetic distances (PD) diversity indices were not significantly different (p=0.14 and 0.15 respectively, Wilcoxon test; Figure 1A). Beta diversity was also assessed through principal component analysis (PCA) using the Euclidean matrix distance (Figure 1B). Statistical permutational analysis of variance (PERMANOVA) showed a significant difference in beta diversity between microbiomes from NT and ASD donors (p=0.001), which was corroborated with the beta dispersion value (p=0.891), indicating that the microbial community composition was distinct between both groups.

**Figure 1.**
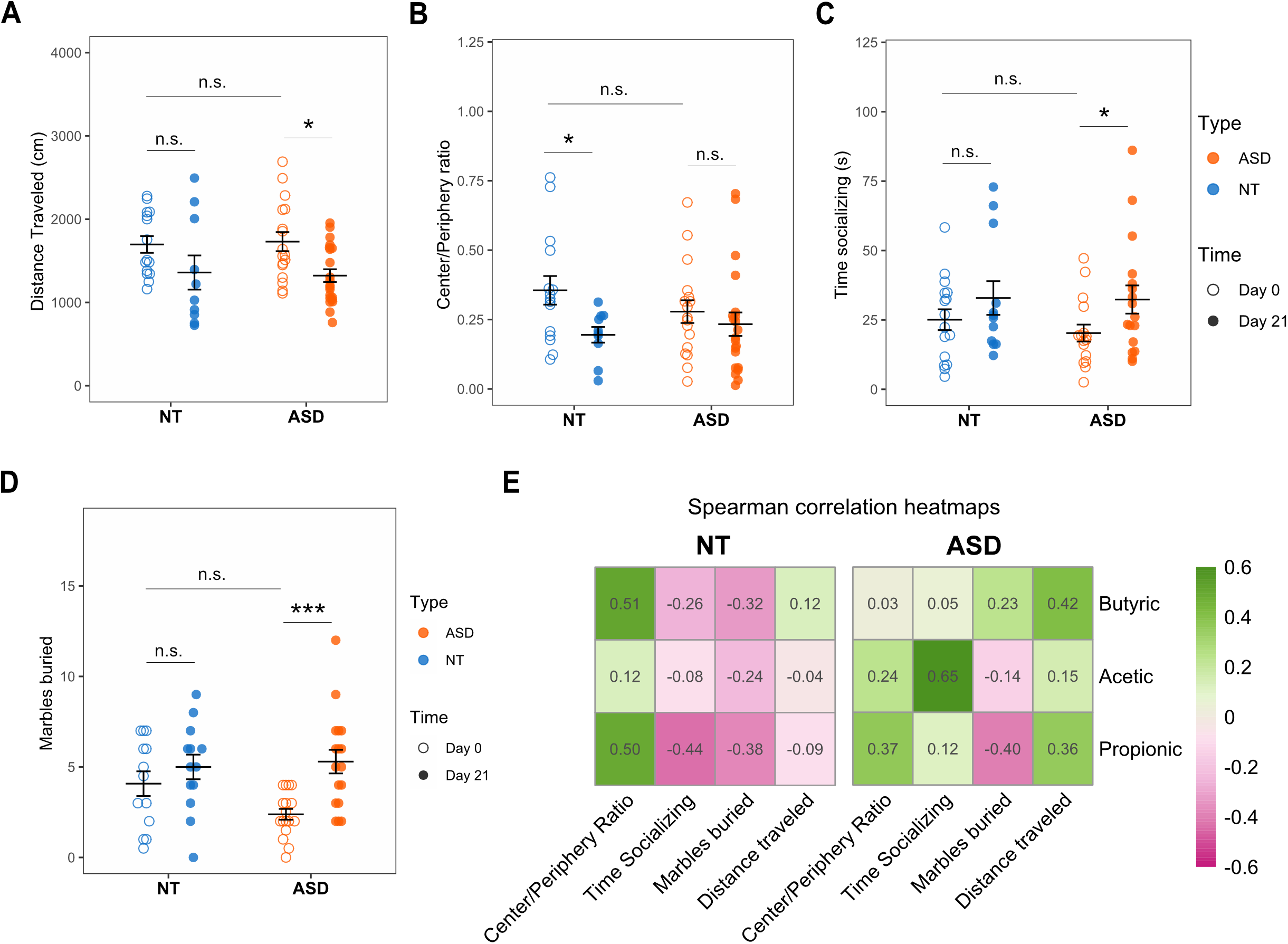
Analysis of gut microbiome in Colombian pediatric patients diagnosed with ASD and NT peers. Bacterial DNA was extracted from fecal samples collected from 7 ASD patients (orange symbols) and 18 NT donors (blue symbols) and analyzed by 16S rRNA sequencing. (A) Analysis of alpha diversity through the observed features, Chao1, Shannon and phylogenetic distances (PD) indices. Data in all box plots are presented as mean and error bars represent outliers. (*) indicates statistically significant differences (p<0.05), “n.s.” denotes no statistically significant differences. (B) PCA plot of Bray Curtis distances, with clustering according to donor status (NT or ASD). Closed symbols depict samples that were used as donors to humanize mice. (C) Differential analysis of ASVs showing specific bacterial genera that were increased in ASD or NT derived microbiota samples

Differential analysis of phylogenetic composition identified 45 statistically different amplicon sequence variants (ASVs) between NT and ASD microbiomes. However, only 30 of those ASVs had a similar percentage above 98.7% with cultivable microorganisms already described, corresponding to 28 distinct genera (Figure 1C). Abundance of *Bacteroides, Bifidobacterium, Blautia, Butyrococcus,* and *Ruminococcus* genera was significantly lower in ASD samples compared to NT counterparts. On the other hand, ASD samples were enriched in ASVs from *Butyricimonas, Collinisella, Dorea, Holdemanella, Phascolarchromobacterium, Sarcina,* and *Prevotella7* and *9* genera.

Taken together, these results show that the gut microbiota of Colombian children diagnosed with ASD is distinct from that of NT children from the same region. In addition, our analysis identified several microbial groups that were differentially modulated in ASD samples and had not been reported in other geographical regions.

### 3.2 Successful humanization of germ-free mice with gut microbiota from donors with and without ASD

We selected specific samples from each group as “donors” to humanize germ-free mice. In total, 4 sample donors were used from either the NT or ASD group of donors, ensuring that they accurately represented the microbial composition of each of these populations (Figure 1B). We obtained male WT mice from the University of Florida Germ-Free Facility (referred to as GF mice), and immediately upon arrival, they were inoculated by oral gavage with fecal slurries prepared from the selected donor samples (3-5 mice per donor; Figure 2A). We confirmed that mice were indeed GF by quantifying the bacterial DNA present in their stool when they arrived from the GF breeding facility and then one week after inoculation. Our quantification showed a significant increase (300-800-fold) in bacterial DNA one week after inoculation (Supplementary Figure 1, p<0.01, Kruskal-Wallis Test).

**Figure 2.**
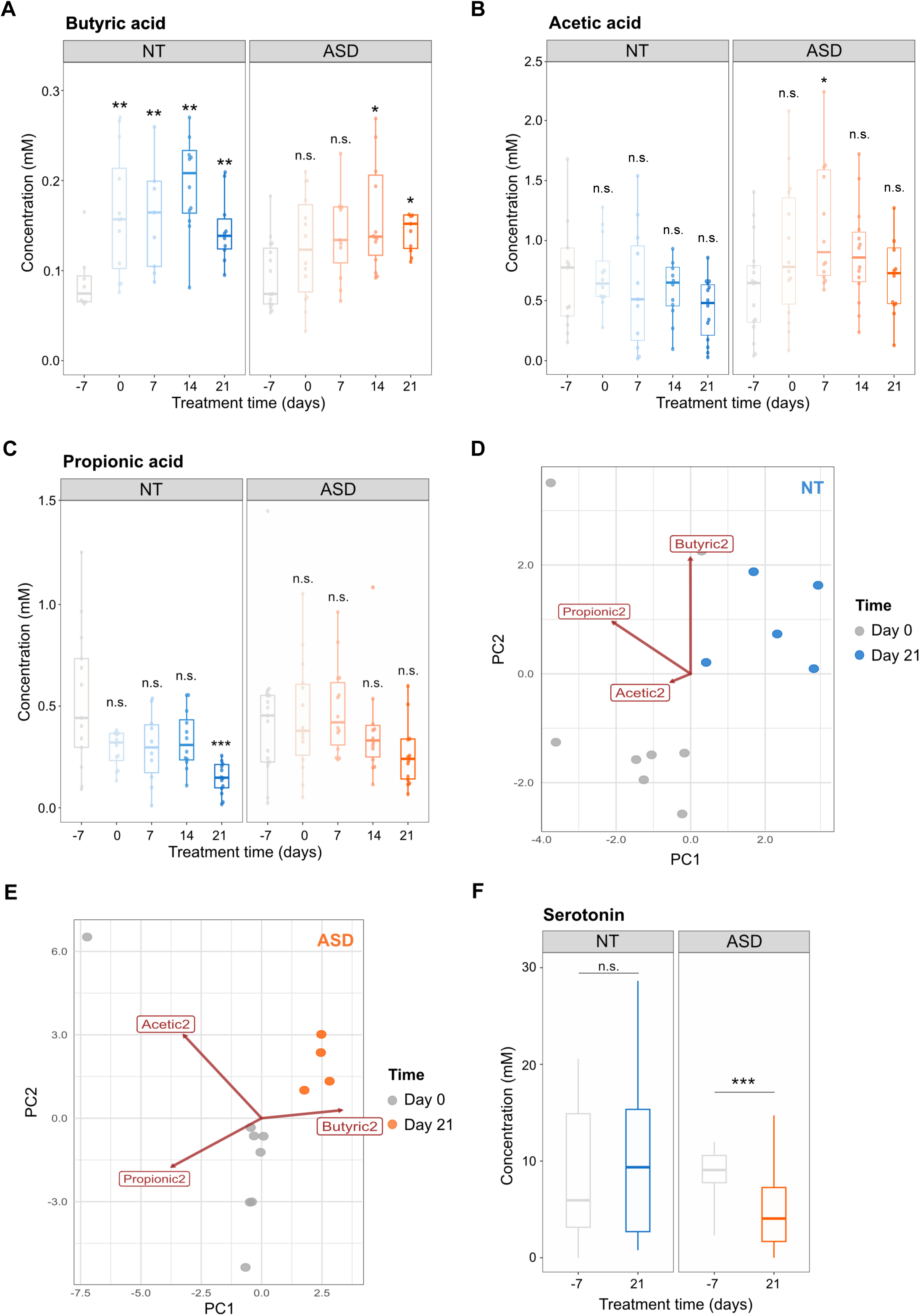
Gut microbiota from Colombian pediatric patients successfully colonizes transplanted GF mice. Four sample donors from NT and ASD groups were used to inoculate 3-5 GF male mice per donor. Fecal samples were then collected to analyze the colonization effectiveness. **(A)** Schematic showing the timeline and experimental plan for GF mice colonization and subsequent microbiota and behavioral intervention. Throughout the figure, blue and orange represent NT and ASD derived samples, respectively. **(B)** Distribution of ASVs found in gut microbiota from colonized mice 4 weeks after transplantation according to origin: corresponding donor, from other donors, and only found in each mouse. **(C)** Relative abundance of ASVs found in gut microbiota from colonized mice according to origin. Each bar represents an individual mouse and bars grouped together depict the same donor (NT 2 to 4, or ASD1 to 4). **(D)** Analysis of alpha diversity (Chao1 and Shannon indices) of gut microbiota at different stages of the humanization process. Data in all box plots are presented as mean and error bars represent outliers. (*) indicates statistically significant differences (*p<0.05 and ***p<0.001), “n.s.” denotes no statistically significant differences. **(E)** PCA plot of Bray Curtis distances for the gut microbiota 4 weeks after colonization, with clustering according to donor status (NT or ASD). **(F)** Comparison (Log2 fold change plot) of differentially regulated bacterial genera between the gut microbiota of mice colonized with NT- or ASD-derived microbiota. Fold-changes were calculated relative to the NT group.

Four weeks post inoculation, we assessed the effectiveness of the gut colonization process (Figure 2B-C). The majority of ASVs found in the colonized GF mice belonged to the corresponding donor (∼59% in NT and ∼52% in ASD samples; Figure 2B). Moreover, these donor-derived microorganisms represented between 75 and 99% of the total relative abundance of gut microbiota present in the colonized mice in both groups (Figure 2C). The results also showed that the colonization efficiency was similar between different donors of the same group (ASD vs. NT; Figure 2B-C). Overall, these results demonstrate that gut microbiota from pediatric Colombian donors successfully colonized GF mice regardless of donor diagnosis.

Further analysis of the samples used for inoculation showed that their alpha diversity decreased significantly from the original fecal donor samples to the slurry prepared for inoculation to the samples collected from colonized mice, as expected for gnotobiotic humanization experiments (Figure 2D, p<0.05, Dunńs Test). Nonetheless, PCA of the Bray-Curtis ordination of gut microbiota after colonization showed that the samples still clustered according to their donor ASD or NT status (Figure 2E; PERMANOVA, p<0.05, Beta-dispersity p>0.05). Furthermore, differential abundance analysis revealed that differences in specific bacterial genera in recipient mice mirrored those observed in the donor population (Figure 1). Specifically, *Prevotella9*, *Butyricicoccus* and *Holdemanella* were significantly increased, while *Bifidobacterium, Faecalimbacterium* and *Akkermansia* were significantly decreased in mice transplanted with microbiota from ASD donors compared to NT counterparts (Figure 2F). Thus, our analyses indicate that the gut microbiota from colonized GF mice represent two distinct populations that capture compositional features of the original donor microbiota.

### 3.3 Treatment with *L. fermentum* K73 synbiotic does not negatively affect the overall health of wild-type mice

Before treating colonized GF mice with the encapsulated *L. fermentum* K73 synbiotic, we assessed its safety profile and modulatory effect on gut microbiota of wildtype (WT) mice (i.e., neither germ-free nor transplanted with human-associated microbiota). Simultaneously, we determined its optimal *in vivo* delivery route by testing two different administration methods: synbiotic incorporated into bread-like food (replacing standard chow) or fed as a powder coating the regular standard chow. WT male mice were divided in 5 groups according to the treatment received: (1) standard chow only (Std) – the control group, (2) carrier-only for the bread-like food (CarBread, see Methods for details of formulation), (3) synbiotic incorporated into the same bread-like food as Group 2 (SynBread), (4) carrier-only in powdered form (CarPowder), and (5) synbiotic in powdered form coating standard chow (SynPowder). The encapsulated *L. fermentum* K73 synbiotic was prepared as previously described (Bauer Estrada, Cardoso Cardenas, et al., 2025). The carrier-only designation refers to the synbiotic encapsulation material composed of whey and high oleic palm oil without *L. fermentum* K73. Both the synbiotic and carrier-only treatments were added by heavily coating the surface of sterile standard chow pellets (10% w/w) while ensuring that total caloric intake remained the same compared to the Std group.

After three weeks of synbiotic treatment, WT mice did not show significant changes in body weight between treatment groups (Supplementary Figure 2A), and all groups consumed comparable amounts of food (daily average ranged between 3.2 to 3.7g, p>0.05, ANOVA; Supplementary Figure 2B). Furthermore, no signs of health issues were observed, and histological analysis showed healthy colons after synbiotic treatment in all groups (Supplementary Figure 2C). Therefore, these findings indicate that the synbiotic formulation is safe to administer to mice and has no detrimental impact on host health *in vivo*.

To investigate possible changes in the gut microbiota, fecal samples were collected before and after synbiotic treatment and then processed for 16S rRNA sequencing. There was no significant difference in alpha diversity between mice grouped in different cages before any treatment (p<0.05, Dunn Test; Figure 3A). We then compared samples from the same mice between Day 0 and Day 21 of treatment with the synbiotic (or its carrier) and observed no significant effect on alpha diversity over time (Figure 3A). When analyzing beta diversity, the community structure of each mouse did not change significantly before and after treatment (Figure 3B, p=0.61, PERMANOVA). Thus, any differences in beta diversity could be attributed to the initial variability among different mouse cages at Day 0 rather than to the effect of the synbiotic treatment. Furthermore, there were no statistically significant differences in relative abundance of phyla and families between different treatment groups (Supplementary Figure 3A-B, p=0.97, ANOVA).

**Figure 3.**
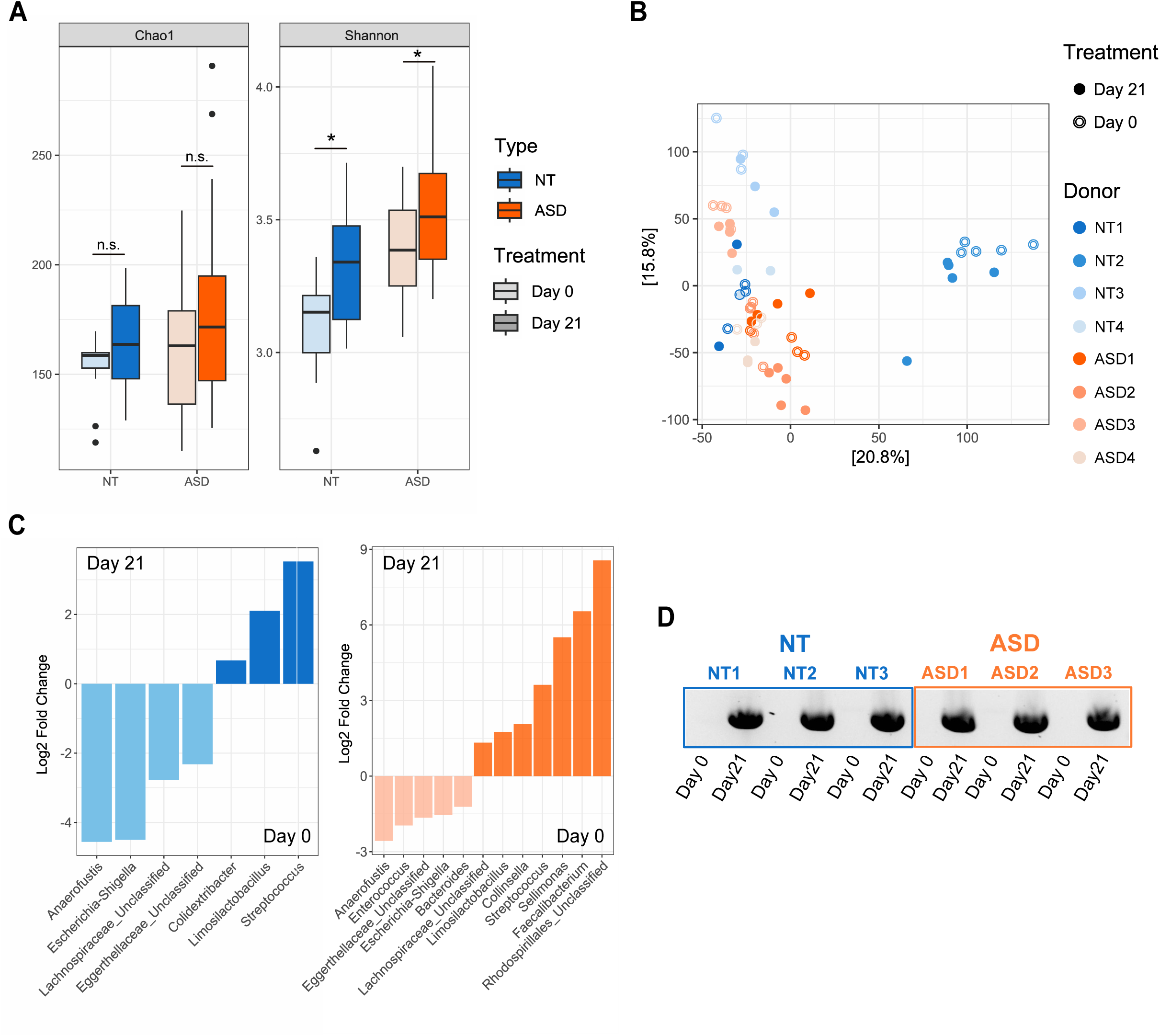
Treatment with *L. fermentum* K73 synbiotic does not alter gut microbiome in wildtype mice. Different groups of wild type (WT) mice (n=5 per group) were fed for three weeks with either standard chow (Std) or our synbiotic in two different presentations, as bread-like food (CarBread for carrier only or SynBread for synbiotic plus carrier), or as powder coating regular chow pellets (CarPowder for carrier only or SynPowder for carrier plus synbiotic). Fecal samples were collected before and after treatment for gut microbiome analysis. **(A)** Analysis of alpha diversity by Chao1 or Shannon indices (left and right panels, respectively) before and after synbiotic treatment. Data in all box plots are presented as mean and error bars represent outliers. (*) indicates statistically significant differences (p<0.05), “n.s.” denotes no statistically significant differences. **(B)** PCA plot of beta diversity, depicting bacterial community structure before (open symbols) and after (colored symbols) synbiotic treatment for all groups. **(C)** Differential analysis of amplicon sequence variants (ASVs) comparing each treatment group to the Std control group.

Differential analysis of ASVs revealed that, as expected, synbiotic treatment promoted the abundance of the *Limosilactobacillus* genus compared to both the Std and corresponding carrier treatment groups (Figure 3C), as confirmed via PCR (Supplementary Figure 3C). Additionally, all treatments increased the relative abundance of the *Ileubacterium* genus while decreasing the abundance of *Rikenella* compared to Std alone. Both genera have been associated with health status in mouse models, with *Ileubacterium* linked to beneficial effects and *Rikenella* to pathogenic effects in some cases (Cox et al., 2017; Gryaznova et al., 2022; Guo et al., 2022; Wu et al., 2024). Our data also showed differences in microbiota modulation based on the synbiotic delivery method (Figure 3C). Compared to the Std-fed control group, administration of the synbiotic through bread decreased the relative abundance of a higher number of taxa than the powdered synbiotic. Moreover, treatment with powdered synbiotic increased the relative abundance of more beneficial microorganisms compared to bread-based synbiotic. Such beneficial groups included the *Enterohabdus* genus and Clostridium UGC002 group, which are known to be butyric acid producers (Samulėnaitė et al., 2024; Zhu et al., 2023).

Collectively, our results demonstrate that both delivery routes are safe for *in vivo* applications of *L. fermentum* K73 synbiotic. Furthermore, the powdered formulation (SynPowder) enhanced taxa that produce beneficial SCFAs, in agreement with our previous *in vitro* and encapsulation studies (Bauer Estrada, Conde-Martínez, et al., 2025). Additionally, SynPowder’s direct application to standard chow eliminates confounding elements from bread-based formulations, offering a streamlined, scalable approach that strengthens its viability for future human supplementation trials while maintaining probiotic efficacy through gastrointestinal transit.

### 3.4 Synbiotic supplementation modifies gut microbiota composition in mice transplanted with human-associated microbiota

After confirming successful GF mice colonization and optimal synbiotic delivery route, we implemented a dietary intervention to determine the effects of the synbiotic in modulating gut dysbiosis and behaviors related to ASD. The encapsulated *L. fermentum* K73 synbiotic was delivered in powder form by coating standard chow pellets and fed to all colonized mice (both NT and ASD groups) for 21 days. All mice readily consumed synbiotic-covered food without significant changes in body weight during treatment (Supplementary Figure 4A), suggesting that the inclusion of the encapsulated synbiotic did not cause food aversion or novelty-related stress (Francois et al., 2022). Additionally, total food intake was comparable between mice transplanted with either NT or ASD associated microbiota (Supplementary Figure 4B), resulting in a synbiotic intake of 0.3±0.03 g/day for both groups. Lastly, synbiotic treatment did not affect the colon integrity and architecture of either group (Supplementary Figure 4C).

Fecal samples were collected weekly to progressively monitor changes in gut microbiota composition during synbiotic treatment. Alpha diversity increased significantly after 21 days of synbiotic treatment in both groups (Figure 4A, Chao1 index for ASD group p=0.34 and for NT p=0.19, Kruskal-Wallis test; Shannon index for ASD group p=0.048 and for NT p=0.014, Wilcoxon test). However, when analyzing beta diversity, we did not observe a global effect of synbiotic treatment on microbiota composition, as the intersubject variability in each donor group was higher than the treatment effect (Figure 4B), in agreement with our previous observations using an *in vitro* digestion system and the same synbiotic and donor-derived microbiota samples (Bauer Estrada, Conde-Martínez, et al., 2025). For this reason, we also generated individual PCA plots for mice colonized with the same donor before (Day 0) and after (Day 21) synbiotic treatment (Supplementary Figure 5). Regardless of donor status (NT or ASD), we consistently observed distinct clustering of samples before and after synbiotic treatment, indicating that all mice responded to the synbiotic treatment irrespective of high intersubject variability.

**Figure 4.**
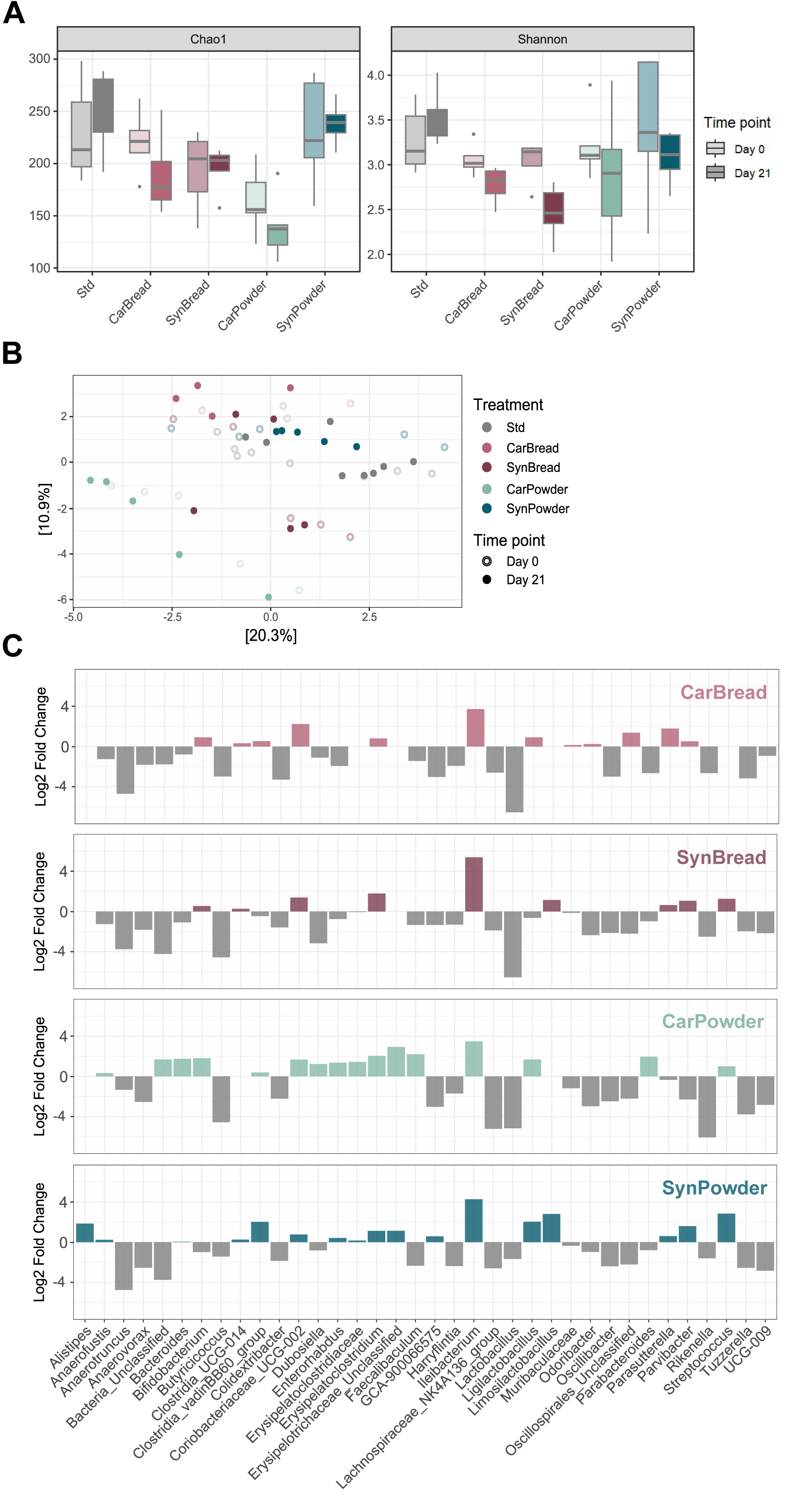
Synbiotic supplementation with *L. fermentum* K73 modifies gut microbiota composition in colonized GF mice. GF mice transplanted with microbiota from either NT or ASD donors were fed for 21 days with synbiotic containing *L. fermentum* K73. Fecal samples were collected before and after treatment, and gut microbiota composition was analyzed. **(A)** Analysis of alpha diversity showing Chao1 and Shannon indices before (lighter color) and after (darker color) synbiotic treatment. Throughout the figure, blue and orange represent NT and ASD derived samples, respectively. (*) indicates significant differences (p<0.05), “n.s.” denotes not significant differences. **(B)** PCA plot of microbiota samples before (open symbols) and after (colored symbols) synbiotic treatment. **(C)** Differential analysis of ASVs from mice transplanted with NT (left panel, in blue) or ASD (right panel, in orange) derived fecal samples. **(D)** PCR analysis confirming the presence of the bile salts hydrolase enzyme gene from *L. fermentum* K73 in colonized mice with NT (left) or ASD (right) derived microbiota. The representative image shows analysis of 3 mice randomly selected per group (M1 to 3), before and after colonization.

We also performed a differential abundance analysis of observed ASVs in fecal samples after 21 days of synbiotic treatment and found several specific ASVs differentially regulated in mice transplanted with either NT or ASD-derived donor microbiota (Figure 4). For instance, synbiotic treatment significantly decreased the relative abundance of *Escherichia-Shigella Eggerthellaceae* and *Lachnospiracea* genera in all mice, regardless of NT or ASD donor status (LM, Wald Test, p<0.05 between day 0 and 21). Additionally, the synbiotic treatment significantly reduced the relative abundance of *Enterococcus and Bacteroides* genera only in mice colonized with ASD-associated microbiota (LM, Wald Test, p<0.05, between day 0 and 21). Interestingly, several bacterial groups showing reduction after synbiotic treatment are known to be pathogenic, suggesting a possible protective effect (Baltazar-Díaz et al., 2022). We also observed an increase in the abundance of *Faecaelibacterium, Sellimonas and Collinsela* (LM, Wald Test, p<0.05, between day 0 and 21), which are known to be butyrate producers and have been shown to be important for gut homeostasis (Muñoz et al., 2020; Parsaei et al., 2021; Zou et al., 2021; Kwon et al., 2023). Lastly, and as expected, we observed a significant increase in ASVs representing *L. fermentum,* which was confirmed by the presence of the bile salts hydrolase enzyme gene from *L. fermentum* K73, as quantified by PCR (Figure 4D). Together, our results show a significant effect of treatment with *L. fermentum* synbiotic in modulating gut microbiota in mice transplanted with human-associated microbiota.

### 3.5 Synbiotic treatment improves the metabolic function of the gut microbiome in mice transplanted with ASD-associated microbiota

SCFAs are key metabolites produced by the gut microbiota that play important roles in energy regulation, immune function, gut barrier integrity and metabolic health among others (Silva et al., 2020; L. Wang et al., 2024). Importantly, SCFAs production has been shown to be dysregulated in ASD patients (Finegold et al., 2010; De Angelis et al., 2013; He et al., 2023; Lagod & Naser, 2023). Thus, we investigated the production of butyric, acetic and propionic acid in mice inoculated with ASD or NT-derived gut microbiota during *L. fermentum* K73 synbiotic treatment (Figure 5, n=12 and 13 mice inoculated with NT and ASD associated microbiota, respectively).

**Figure 5.**
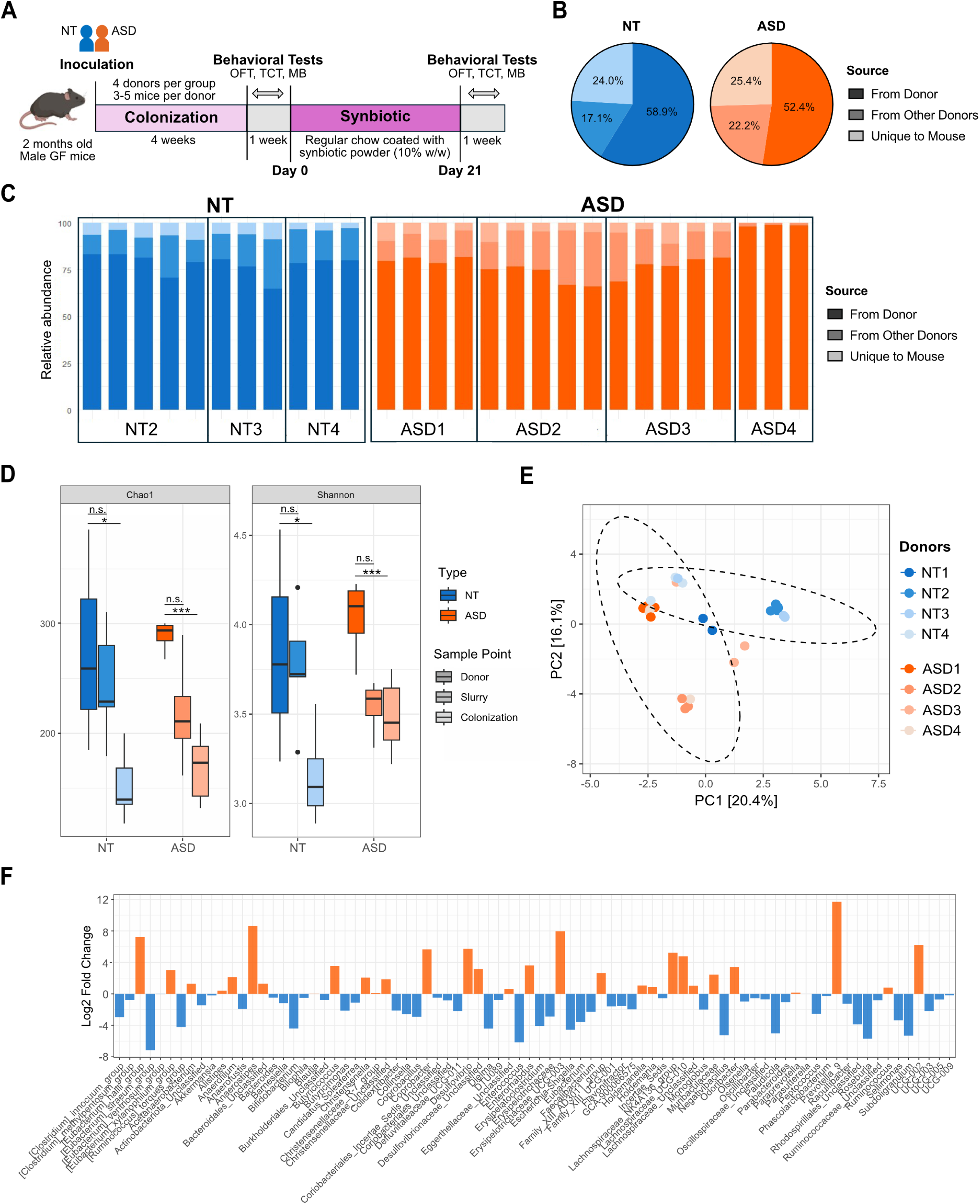
Synbiotic treatment differentially modulates metabolic function in gut microbiome of colonized GF mice. Fecal samples were collected weekly throughout the synbiotic treatment from GF mice that were colonized with either NT or ASD-derived gut microbiota. These fecal samples were then processed to quantify levels of metabolites including several SCFAs and serotonin. **(A-C)** Quantification of fecal concentration of **(A)** butyric acid, **(B)** acetic acid, and **(C)** propionic acid in samples from mice colonized with NT (blue, n=12 mice) or ASD (orange, n=13 mice) derived microbiota. Samples were collected starting before inoculation (Day −7) until week 3 of synbiotic treatment (Day 21). (*) indicates statistically significant differences (*p<0.05, ***p<0.001), “n.s.” denotes no statistically significant differences. **(D)** and **(E)** PCA plots of Bray distances before (Day 0) and after (Day 21) synbiotic treatment for the gut microbiota of NT-(blue symbols) or ASD-(orange symbols) colonized mice. Red arrows show concentration of fecal SCFA species (Butyric, Acetic and Propionic acid) used as constraints. **(F)** Quantification of serotonin in fecal samples from mice colonized with NT-(blue) or ASD-derived (orange) microbiota. (***) indicates statistically significant differences (p<0.001), “n.s.” denotes not statistically different.

Fecal butyric acid levels increased significantly over time in mice colonized with either NT or ASD-associated gut microbiota (Figure 5A; p<0.05, Dunn’s Test). This change correlated with the observed increase in abundance of specific microorganisms that are known producers of butyric acid and have been considered beneficial for human health such as *Sellimonas* and *Faecalibacterium* (Figure 4D-E) (Muñoz et al., 2020; Zou et al., 2021; Martín et al., 2017). While acetic acid levels did not change in mice colonized with NT donor-derived gut microbiota, there was a statistically significant but transient increase in acetic acid production in mice transplanted with ASD donor-derived microbiota (Figure 5B, p<0.05, Dunn’s Test). Finally, levels of propionic acid decreased significantly by day 21 in mice transplanted with NT-derived gut microbiota, with no change in the ASD-inoculated counterparts (Figure 5C; p>0.05, Dunn’s Test). Additional Beta diversity analysis grouping mice by individual donor sample and SCFA production (Figures 5D-E), showed that both types of donor microbiota (NT and ASD), clustered in groups corresponding to before (Day 0) and after (Day 21) synbiotic treatment. Using SCFA concentration as constraints, we observed that synbiotic treatment appeared to promote the production of both butyric and acetic acid, but not propionic acid (Figures 5D-E), in agreement with our prior observations (Bauer Estrada, Conde-Martínez, et al., 2025).

We also quantified the production of serotonin, another important gut microbiota-derived metabolite and neurotransmitter that plays key roles in mood regulation, immune system, cognitive function and intestinal motility (Sanidad et al., 2024; Potter et al., 2024; Nunzi et al., 2025). In addition, elevated whole-blood serotonin levels have been consistently reported in children with ASD (Hanley, 1977; Veenstra-VanderWeele et al., 2012; Abdulamir et al., 2018). After treatment with *L. fermentum* K73 synbiotic, fecal serotonin levels significantly decreased in mice colonized with ASD donor-derived gut microbiota (Figure 5F), from 8.4±3.5 nM/mg before to 5.0±1.4 nM/mg after synbiotic treatment (p<0.01, Kruskal-Wallis test). In contrast, fecal serotonin levels did not change in mice colonized with NT donor-derived gut microbiota in response to synbiotic treatment (average serotonin levels before treatment were 8.9±2.5 and after 10.9±3.2 nM/mg, p=0.66; Kruskal-Wallis test). In addition, quantification of circulating serotonin from blood samples collected at the end of the behavioral analyses showed no significant differences between NT or ASD colonized mice (Supplementary Figure 6, p>0.15; Kruskal-Wallis test).

Taking together, our results show that synbiotic treatment differentially modulates the gut production of specific SCFAs and serotonin in colonized mice. This suggests a potential modulatory effect on metabolic processes that are important for gut microbial community, which, in turn, might be influential for the overall health of the host.

### 3.6 Synbiotic intervention alters social and exploratory behavior in mice colonized with ASD derived microbiota

Because ASD patients display several key behavioral changes, including altered social behavior, hyperactivity and repetitive behaviors, amongst others (American Psychiatric Association, 2013), we set out to determine the potential behavioral effects of treatment with encapsulated *L. fermentum* synbiotic on colonized mice. We specifically focused on well-established mouse tests to investigate social and exploratory behaviors that have been shown to be altered in ASD mouse models (Kazdoba et al., 2016). These tests include the open field test, three chamber test, and marble burying test. Importantly, to document progressive changes over time, these behavioral tests were administered to all mice before and after the synbiotic dietary intervention.

Total distance traveled in the open field test (OFT) showed no difference between mice colonized with NT or ASD microbiota before synbiotic treatment (Day 0; Figure 6A). The average distance covered for NT colonized mice was 1,696±389cm before treatment, while ASD colonized mice moved an average of 1,730±473cm (n=15 and 17 mice for NT and ASD groups, respectively, for all behavioral experiments; p=0.81, Kruskal-Wallis test). Moreover, only mice transplanted with ASD-derived microbiota showed a significant reduction in total exploration after synbiotic treatment. The average distance covered for NT colonized mice was 1,359±649cm after synbiotic treatment, while ASD colonized moved an average of 1,322±352cm (Figure 6A, p=0.09 and p<0.05 for NT and ASD colonized groups, respectively).

**Figure 6.**
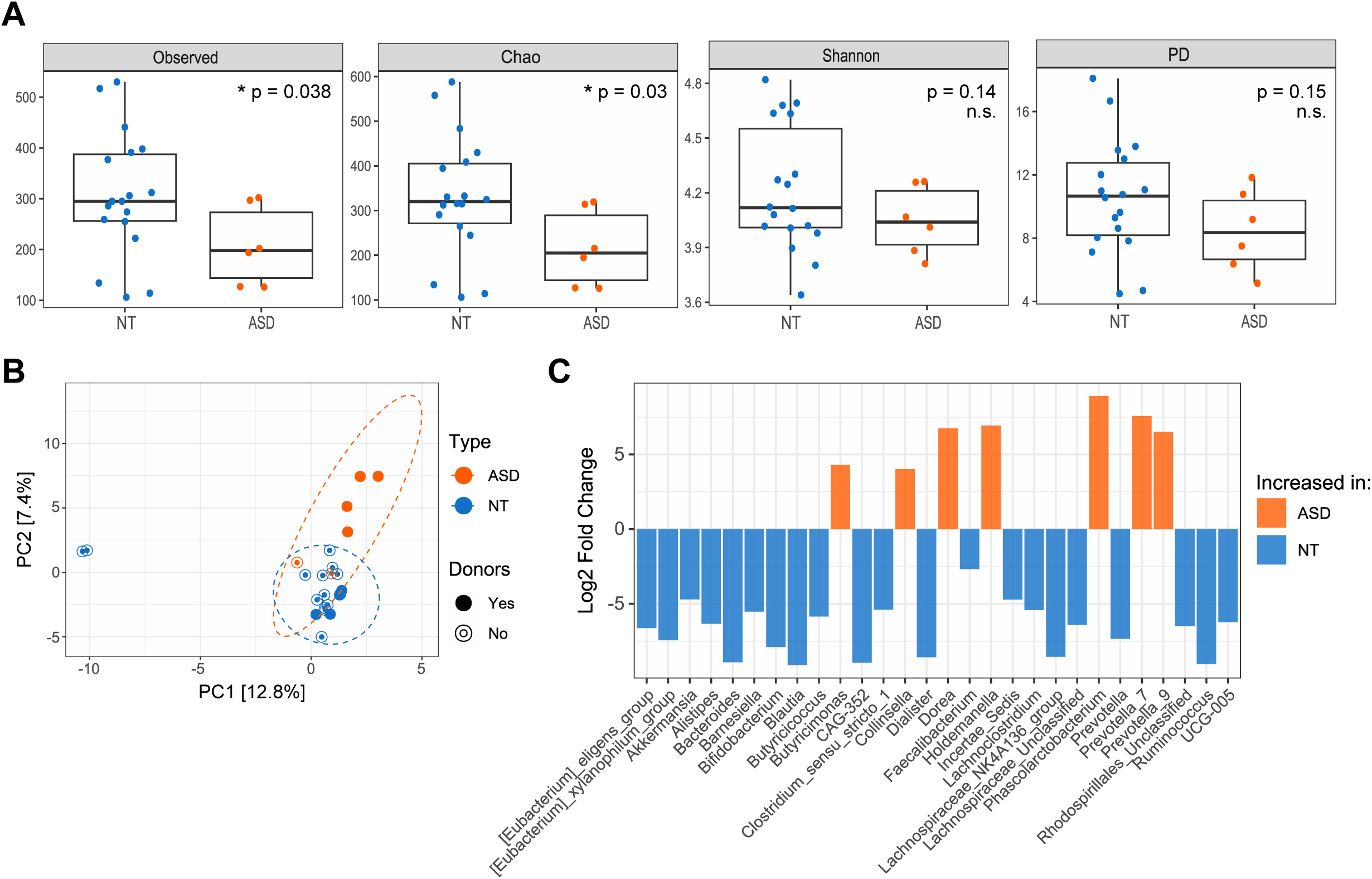
Synbiotic intervention differentially alters social/exploratory behaviors in mice colonized with NT or ASD-derived microbiota. GF mice transplanted with NT (blue symbols, n=12 mice) or ASD (orange symbols, n=13 mice)-derived microbiota were administered synbiotic for 21 days. After this treatment, all mice were subjected to a battery of behavioral tests to investigate associated behavioral outcomes: **(A)** total distance covered, **(B)** time spent in the center vs periphery in the open field test (OFT), **(C)** time socializing in the three-chamber test (TCT), and **(D)** quantification of performance in the marble burying test (MBT) based on the total number of marbles buried. Data are presented as mean ± SEM. (*) indicates statistically significant differences (*p<0.05 and ***p<0.001), “n.s.” denotes not statistically different. **(E)** Spearman correlation analysis between behavioral outcomes and fecal SCFA concentration in mice transplanted with NT (left) or ASD-derived microbiota (right). (*) depicts statistically significant differences (p<0.05).

We then quantified time spent in the center vs the periphery of the OFT, as a proxy for anxiety-like behaviors in rodents. Rodents generally avoid the center of an open field and gravitate towards the periphery, behavior called thigmotaxis (Simon et al., 1994), and a reduction in time spent in the center is indicative of heightened anxiety-like behaviors (Seibenhener & Wooten, 2015b). We observed that before synbiotic treatment (Day 0) there was no difference between mice colonized with NT or ASD microbiota in the center to periphery (C/P) ratio (see Methods). The average C/P ratio for NT colonized mice was 0.36±0.2 before treatment, while that of ASD colonized was 0.28±0.17 (Figure 6B, p=0.25 for Day 0 between NT and ASD group, Kruskal-Wallis test). Interestingly, after synbiotic treatment (Day 21) the C/P ratio was not affected in mice colonized with ASD derived microbiota while decreasing significantly in the NT colonized group. The average C/P ratio for NT and ASD colonized mice was 0.195±0.09 and 0.23±0.19 after synbiotic treatment (Figure 6B, p=0.031 for NT and p=0.25 for ASD groups).

To evaluate social interactions between mice, we used the three-chamber test (TCT). Before synbiotic treatment (Day 0), the average time socializing for NT colonized mice was 25.1±15.0s and 20.3±12.0s for ASD colonized mice (Figure 6C, p=0.39, Kruskal-Wallis test). After the synbiotic treatment (Day 21), however, there was a significant increase in time socializing only in the ASD colonized group. In this case, the average time socializing for NT colonized mice was 32.9±21.0s, while ASD colonized mice spent an average of 32.3±20.0s with a cage mate after synbiotic treatment (p=0.52 and p<0.05 for NT and ASD groups respectively). This is consistent with previous literature showing that ASD mouse models do not always display reduced preference for a social stimulus in baseline conditions (Rein et al., 2020). Nonetheless, the synbiotic treatment increased social behavior of mice colonized with ASD derived microbiota, with no change in the NT colonized group.

Lastly, before the synbiotic treatment (Day 0), while mice transplanted with both NT- and ASD-associated microbiota showed comparable behavior in the marble burying test (MBT), there was a clear trend towards reduced performance in ASD colonized mice. The average number of buried marbles for NT colonized mice was 4.1±2.5 before treatment, compared to 2.4±1.2 marbles in ASD colonized mice (Figure 6D, p=0.06, Kruskal-Wallis test). Importantly, after synbiotic treatment (Day 21), only mice colonized with microbiota from ASD donors showed an increase in MBT performance. The average number of buried marbles for NT colonized mice was 5.0±2.5 after treatment (p=0.42 comparing before and after synbiotic), while ASD colonized mice buried an average of 5.3±2.7 marbles on Day 21, doubling the amount from before synbiotic supplementation (p<0.001, Kruskal-Wallis test, comparing before and after synbiotic).These findings align with previous studies of humanized mouse models and GF mice, showing reduced MBT performance compared to WT counterparts (Park et al., 2025). Our results showing that synbiotic treatment led to an increase in marbles buried only in ASD colonized mice suggest a normalizing effect in this behavioral task.

To gain a better understanding of how the observed changes in gut microbiota composition might relate to behavioral outcomes, we conducted a Spearman correlation analysis including quantified behavior variables and SCFAs production after synbiotic treatment (Figure 6E). Overall, this analysis showed that some associations between specific SCFAs and behavioral changes were different between NT and ASD colonized mice. For instance, acetic acid production had a strongly positive correlation (r=0.65, p<0.05) with sociability in mice that were transplanted with ASD derived microbiota, while no correlation (r=-0.08, p>0.05) was observed in mice colonized with NT derived microbiota. Also, butyric acid was positively correlated with OFT C/P ratio in mice transplanted with microbiota from NT donors (r=0.51, p>0.05), while mice transplanted with ASD microbiota showed no association (r=0.03, p>0.05). On the contrary, other behaviors were equally modulated by the same SCFA regardless of the NT or ASD status of the microbiota donor. For example, marble burying behavior was negatively correlated with propionic acid for both NT-(r=-0.38, p>0.05) and ASD-colonized mice (r=-0.40, p>0.05), indicating a possible association between propionate production and repetitive behaviors (MacFabe et al., 2011).

Overall, our behavioral analysis showed that dietary intervention with *L. fermentum* K73 synbiotic resulted in modulation of several behaviors that have been described as negatively changed in ASD mouse models. Notably, these changes were observed primarily in mice transplanted with ASD-derived microbiota, supporting the idea that this synbiotic could be an effective therapeutic intervention to restore gut microbiota function. Our analysis of the correlation of specific SCFAs and behavioral outcomes further suggests that gut microbiota metabolism might be an important factor to consider when anticipating the effects of synbiotic supplementation.

## 4. DISCUSSION

Synbiotic formulations that combine probiotics and prebiotics are promising dietary tools to restore gut microbial balance and modulate the microbiota–gut–brain axis (Chung et al., 2016; Ristori et al., 2019). Current attempts to treat neurodevelopmental disorders (including ASD) with broadly applied probiotic mixtures have yielded limited benefits, in part because they overlook the ecological complexity of the human gut microbiome, which is shaped by host genetics, diet, and environmental context (Lu et al., 2025; Zhou et al., 2025). This limitation is particularly acute in Latin America, where rapid nutritional transitions, socio-economic inequalities, and extensive genetic admixture intersect with a growing burden of ASD, even as most microbiome research remains concentrated in high-income countries (W. Liu et al., 2025). Consequently, major global microbiome initiatives continue to omit Latin American populations, leaving much of the Global South underrepresented in current microbiome science (Shao et al., 2026). Within this context, we show that a *L. fermentum* K73 synbiotic modulated human-derived gut microbiota in gnotobiotic mice colonized with microbiota from Colombian children with and without ASD, increasing beneficial taxa, altering key SCFAs, and associating with changes in ASD-relevant behaviors. To our knowledge, our study documents the first successful colonization of GF mice with Colombian pediatric microbiota, supporting targeted synbiotics as a potential strategy to improve gut microbiome function in health and disease.

### 4.1 Synbiotics as effective modulators of gut microbiota composition and function

*L. fermentum* K73 synbiotic was well tolerated, readily consumed, and shifted the composition and functional activity of human-derived gut microbiota in colonized GF mice. Furthermore, synbiotic treatment decreased taxa associated with pathogenicity or dysbiosis in ASD including *B. thetaiotamicron*, *Shigella,* and *Escherichia* (Carmel et al., 2023; D. Wang et al., 2025), and increased taxa linked to beneficial outcomes in ASD and other neurological conditions, such as *Faecalibacterium prausnitzii*, *Sellimonas*, and *Streptoccus termophilus* (Parsaei et al., 2021; Sokol et al., 2008; Coretti et al., 2018; Kim et al., 2022; Y. Zhang et al., 2024). Stronger shifts in these taxa were observed in ASD-derived communities, suggesting that dysbiotic microbiota may be particularly responsive to synbiotic intervention.

Synbiotic supplementation also altered SCFA production, increasing butyrate-producers like *F. prausnitzii* (Martín et al., 2017; Parsaei et al., 2021) in both NT- and ASD-derived microbiota and transiently increasing acetate in ASD-colonized mice, consistent with the enrichment of known SCFA producers and the roles of these metabolites in maintaining gut barrier integrity, metabolic regulation, and neuroimmune modulation (Nagpal et al., 2018; Tom Dieck et al., 2021; Singh et al., 2022; Q. Zhang et al., 2024). Microbial acetate has been linked with metabolic health benefits, including lower fasting blood glucose and improved lipid metabolism, and the reduction of inflammation, contributing to improved mood and cognitive outcomes (Osman et al., 2023). However, this increase in acetic acid was transient, differing from prior work showing that supplementation with a mixture of *Lactobacillus* and *Bifidobacterium* probiotics increased serum and brain levels of SCFAs, including acetate and butyrate, in Alzheimer’s disease mouse models (Kaur et al., 2020; Van Zanten et al., 2012). Responses differed from some prior probiotic studies, likely due to differences in models, donor microbiota, and formulations, underscoring that SCFA dynamics are highly context dependent.

In addition to SCFA changes, fecal serotonin was reduced in mice colonized with ASD donor microbiota following synbiotic supplementation. Because serotonin regulates a variety of important physiological functions, this targeted reduction suggests an additional mechanism through which synbiotics may potentially modulate the ASD-associated gut microbiome–brain axis. The lack of similar effect in mice colonized with microbiota from NT donors further indicates a context-dependent microbiota-modulating action rather than a nonspecific suppression of serotonin synthesis. Moreover, fecal serotonin can arise from several gut associated sources, including host enterochromaffin cells located in the gastrointestinal tract (the primary source), direct microbial synthesis and dietary components such as tryptophan and SCFAs (Banskota & Khan, 2022). Importantly, our study did not directly test causal links between synbiotic supplementation and gut-derived serotonin, nor can our findings distinguish which of these serotonin sources were specifically affected.

A notable result was the strong agreement between these *in vivo* findings and prior *in vitro* digestion and batch fermentation assays performed with the same donors and synbiotic (Bauer Estrada, Conde-Martínez, et al., 2025), including parallel changes in specific taxa and SCFA profiles. This cross-platform consistency supports donor-matched *in vitro* bioreactors as useful early-stage screening tools for predicting microbial and metabolic responses to synbiotic interventions and complements current efforts to expand scalable, non-animal approaches for microbiome-targeted therapeutic development (C. Liu et al., 2023; Clavel et al., 2025). Furthermore, despite key limitations such as the small number of human donors and the limited set of transplanted communities, this pilot study demonstrates feasibility, successful colonization, safety, and preliminary efficacy, indicating that the *L. fermentum* K73 synbiotic can modulate microbiota function and behavior in a humanized mouse model and providing a foundation for larger, more diverse studies aimed at clinical translation.

### 4.2 Association between gut dysbiosis and neurodevelopmental disorders

Our findings contribute to a growing body of evidence supporting an association between gut dysbiosis and neurodevelopmental disorders, including ASD, and provide new microbiome data from an understudied Latin American population (Almeida et al., 2019; Pasolli et al., 2019; Abdill et al., 2022). While our study did not compare gut microbiome profiles from different geographical regions, in fact, all samples came from donors residing in the same city (Bogotá, Colombia), ASD-associated microbiota showed reduced richness and distinct community structures compared to NT peers, aligning with studies from other geographical areas(Ding et al., 2021; Porras et al., 2021; Pulikkan et al., 2018; Zurita et al., 2020). Our results also showed differences in major groups such as Proteobacteria and Bacteroides species when compared with distinct cohorts from North America, Europe, and Asia (Finegold et al., 2010; Ding et al., 2021; Williams et al., 2011; Vernocchi et al., 2022; Aziz-Zadeh et al., 2025).

We also identified taxa not previously highlighted in the context of ASD, including reduced fiber-degrading genera (Mukherjee et al., 2020) (for example, *Barnesiella, Butyricicoccus, Eubacterium, Ruminococcus*) and enrichment of *Collinsella* (Gomez-Arango et al., 2018), and the propionate-producing *Phascolarobacterium* in ASD samples, consistent with lower fiber intake and potential links between propionate and ASD-like features. These results emphasize both shared and population-specific microbial signatures associated with ASD and underline the importance of geographic and dietary context. Larger studies, including regional initiatives such as the LatinBiota Consortium (Reyes et al., 2025), will be needed to define robust ASD-associated microbiome features across the region.

Our results should nonetheless be interpreted considering the disparity in donor group sizes (7 ASD vs 18 NT), which reflects the limited availability of donors meeting our inclusion and exclusion criteria within a specific clinical population. This imbalance is likely to reduce the statistical power to detect subtler differences in richness, community structure, and taxa abundance between groups. Hence, compositional differences may have gone undetected and the magnitude of those reported here should be interpreted with the appropriate caution. The taxonomic and diversity signatures described above are therefore best considered preliminary characterizations of ASD-associated microbial dysbiosis in this Colombian population, to be validated in larger and more balanced cohorts.

### 4.3 Successful colonization of GF mice with human Colombian microbiota

This study represents, to our knowledge, the first successful transplantation of GF mice with gut microbiota from Colombian pediatric donors, extending prior humanization work in Latin American populations (Porras et al., 2021). A subset of donor microorganisms (approximately 20–50% of ASVs) established in mice but accounted for most of the relative abundance (70-90%), and colonization efficiency was similar between NT- and ASD-associated communities. Our observations are consistent with other GF colonization studies (Collins et al., 2015; Gray et al., 2024; Lundberg et al., 2020; Porras et al., 2021) and expected reductions in alpha diversity due to differences in mouse gut environment (Gustafson et al., 2025; Li et al., 2022) and sample preparation procedures (Superdock et al., 2023).

Beta diversity analyses showed clear clustering by donor ASD versus NT status, and ASD-associated genera such as *Collinsella* and *Dorea* were increased in ASD-colonized mice, in line with previous links to ASD severity, SCFA metabolism, and the production of pro-inflammatory cytokines in children with ASD (Morton et al., 2023; Carmel et al., 2023; de Vadder & Mithieux, 2018; F. Liu et al., 2019). Overall, these results demonstrate robust establishment of a colonized mouse model using Colombian microbiota, providing a platform to study gut microbiome–host interactions and to evaluate microbiome-targeted interventions tailored to this understudied population.

### 4.4 Behavioral effects of synbiotic intervention in transplanted mice and correlation with microbiota changes

Several studies have linked gut microbiota composition and function to behavioral outcomes in humans diagnosed with ASD (Christian et al., 2015; Kang et al., 2013) and animal models(Buffington et al., 2021; Hsiao et al., 2013; Xiao et al., 2021). However, studies with colonized GF mice with ASD-derived microbiota have shown mixed results (Gustafson et al., 2025; Luczynski et al., 2016; Roussin et al., 2024). Consistent with these findings, we did not detect strong baseline behavioral differences between ASD- or NT-colonized mice. After synbiotic supplementation, however, ASD-colonized mice exhibited reduced locomotion and anxiety-like behaviors in the open field, increased sociability, and enhanced marble-burying performance toward patterns typical of conventionally raised mice (Roussin et al., 2024; Diaz Heijtz et al., 2011; Desbonnet et al., 2014; Spichak et al., 2018), suggesting that synbiotic-driven modification of dysbiosis can potentially modulate multiple ASD-relevant behavioral domains.

These behavioral changes were accompanied by specific shifts in gut metabolites, including SCFA and serotonin. Increased acetate production correlated with improved sociability in ASD-colonized mice, whereas reduced propionic acid in all colonized mice negatively correlated with marble burying behavior. These observed associations should be interpreted cautiously as this study was not designed to explicitly test causal relationship between microbial metabolites and behavior. Nonetheless, our observations align with a broader literature linking SCFAs to social and exploratory behavior (van de Wouw et al., 2018; Marrocco et al., 2022; T. Wang et al., 2022, 2023). Specifically, acetate has been associated with enhanced social behavior and reduced anxiety in ASD-like mouse models (Osman et al., 2023; McFarlane et al., 2008; Kratsman et al., 2016) and elevated propionate has been shown to promote ASD-like phenotypes (Shultz et al., 2008; Alsubaiei et al., 2022). Similarly, synbiotic treatment was associated with reduced fecal serotonin only in ASD-colonized mice suggesting potential modulation of dysregulated peripheral serotonin, which has been shown to be elevated in ASD patients (Esposito et al., 2024; Israelyan & Margolis, 2019; Muller et al., 2016). However, we did not detect any specific association between serotonin and behavioral outcomes. Because central and peripheral serotonin are regulated by distinct, functionally segregated mechanisms (Muller et al., 2016), and because we did not measure brain serotonin or baseline circulating serotonin prior to synbiotic treatment, we cannot determine how synbiotic-related changes in gut-derived serotonin might relate to central serotonergic function or behavior. Nonetheless, the coordinated changes in SCFAs, serotonin levels, and ASD-like behavior suggest that the gut–brain axis contributes to the response to synbiotic treatment, although causal underlying mechanisms require further study.

## 5. CONCLUSION

In summary, this pilot study shows that supplementation with *L. fermentum* K73 synbiotic can modulate human-derived gut microbiota in a gnotobiotic mouse model. This supplement was particularly effective if that microbiota originated from donors diagnosed with ASD. These microbiota shifts were accompanied by altered SCFA profiles, changes in microbiota-regulated serotonin, and differences in behavioral outcomes, suggesting that synbiotic-driven restoration of dysbiosis can influence multiple host physiological processes along the gut–brain axis. In addition, our work establishes feasibility and safety *in vivo* and highlights the utility of complementary in vitro digestion models for predicting microbiome responses to synbiotic interventions. More broadly, it advances microbiota-targeted strategies in an understudied population and underscores the need for expanded research across diverse geographic and genetic backgrounds to support the development of effective, context-specific therapeutic applications.

## Supporting information

BauerEstrada_et_al.2026Suppl.Figures

## Ethics approval and consent to participate

Collection of fecal samples for this study was approved by The Ethical Committee of the Universidad de La Sabana and Clinica Universidad de La Sabana (Act of session No. 100). The research activities were performed following the Declarations of Helsinki for the Ethical Principles for Medical Research Involving Human Participants. Informed consent was obtained from the legal guardians of all participants. The use of selected fecal samples from human donors to colonize gnotobiotic mice was approved by the University of Florida Institutional Review Board (Protocol #IRB202302069).

## Data Availability

Raw sequencing data from human donors is available in the Bioproject accession number PRJNA1121380. Mouse sequencing data from are being submitted to NCBI and available upon request until they are publicly available. The Bioproject accession number will be obtained prior to acceptance for publication.

## Authors’ contributions

Conceptualization and methodology: KBE, AAG, MXQC, AMP, and KA; investigation (data collection): KBE, KMMA, CW, SDS, KFS, VM, NCM, and KA; formal analysis: KBE, KMMA, CW, SDS, KFS, VM, AAG, NCM, AMP, and KA; visualization: KBE, KMMA, AAG, AMP, and KA; writing – original draft: KBE, AMP, and KA; writing – review & editing: KBE, MXQC, AMP, and KA; supervision and funding acquisition: MXQC, AMP, and KA. All authors approved the final version of the manuscript.

## Acknowledgements

This project was possible thanks to the contract “Contrato Marco de Acceso a Recursos Genéticos y sus Productos Derivados 324 −2021 (RGE0381)” granted by the Colombian Ministry of Environment and Sustainable Development (Ministerio de Ambiente y Desarrollo Sostenible de Colombia). We are also very grateful to the “Apushi Para El Desarrollo de La Niñez” Foundation for their support during this study.

