## Supplementary material for "*Limosilactobacillus fermentum* K73 synbiotic modulates gut microbiota function and behavior in gnotobiotic mice transplanted with microbiota from children diagnosed with autism spectrum disorder": BauerEstrada_et_al.2026Suppl.Figures

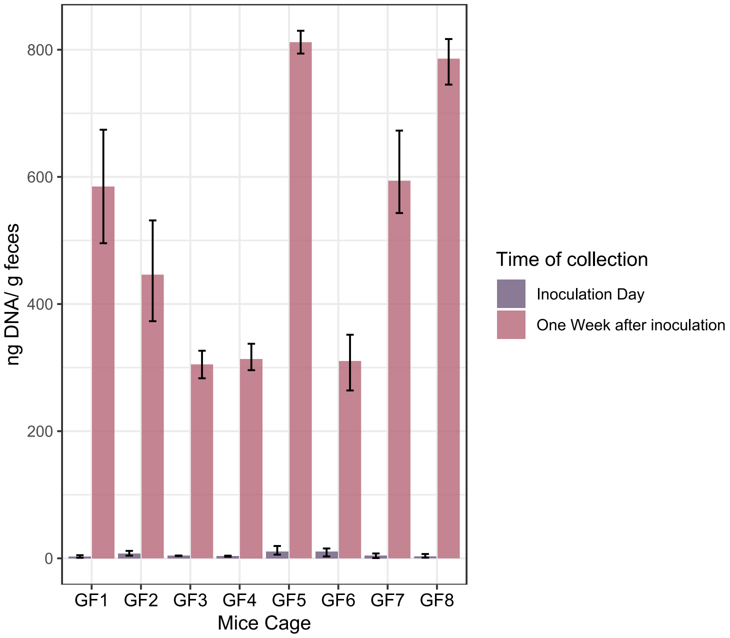


**Supplementary Figure 1. Quantification of bacterial DNA in fecal samples from GF mice confirms germ-free (GF) status and successful inoculation with human microbiota.** Bacterial DNA was extracted and quantified from fecal samples were collected from GF mice at the time of arrival from the GF facilities prior to inoculation (purple bar), and one week after inoculation with human-associated microbiota (dark pink bar). Data are shown as mean ± SD, normalized to sample weight and averaged per mouse cage (n=3 to 5 mice per cage).

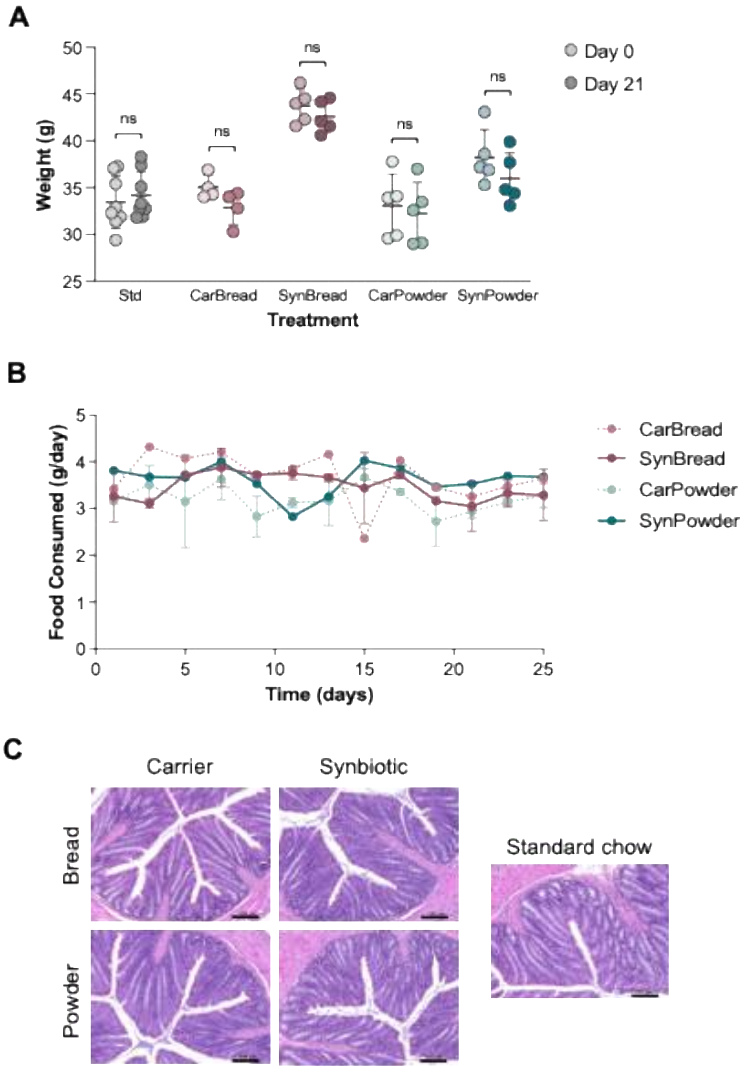


**Supplementary Figure 2. Treatment with carrier or synbiotic formulations does not impact the health status of WT mice.** Different groups of adult male C57Bl/6 WT mice (n=5 per group) were fed for 21 days with either standard chow (Std) or our synbiotic in two different presentations, as bread-like food (CarBread for carrier only or SynBread for synbiotic plus carrier), or as powder coating regular chow pellets (CarPowder for carrier only or SynPowder for carrier plus synbiotic). **(A)** Mouse body weight at days 0 and 21 of treatment. Data are presented as mean ± SEM. "n.s." denotes no significant differences (ANOVA, $p>0.05$). **(B)** Average food consumption per group, measured every other day throughout the course of the experiment, was not statistically different. **(C)** Representative images of H&E-stained colon sections for each experimental group 21 days after treatment. Scale bar represents 100 μm.

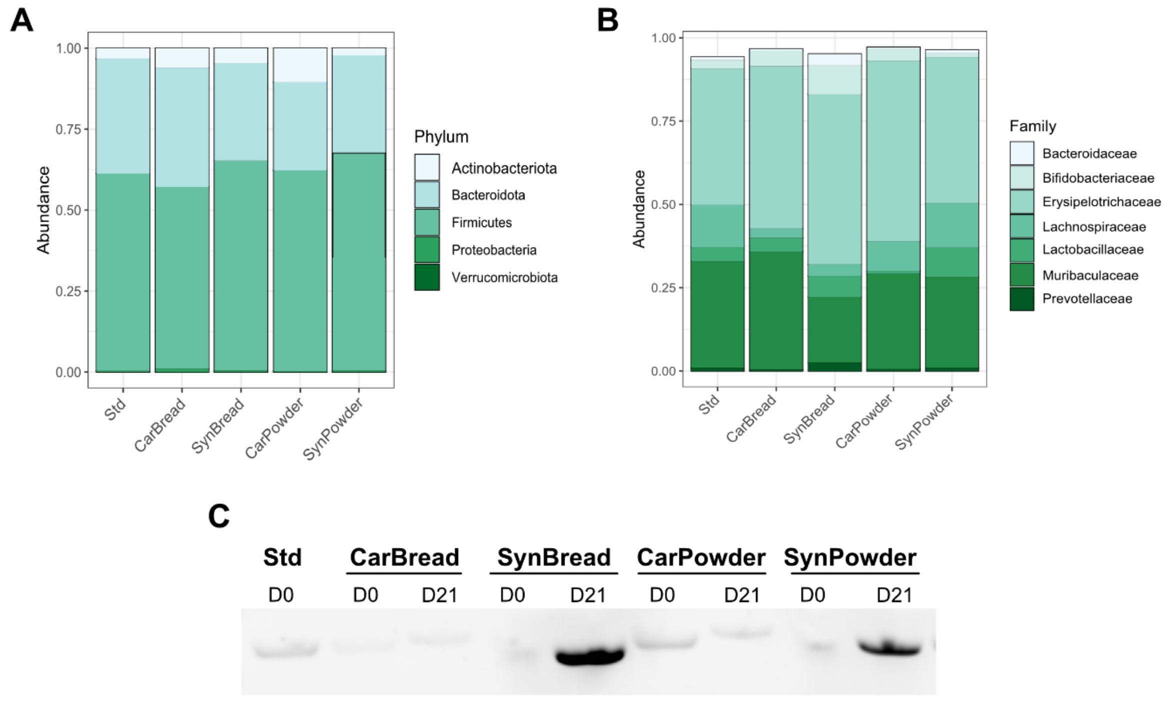


**Supplementary Figure 3. Taxonomic analysis of gut microbiota composition in WT mice after treatment with carrier and synbiotic formulations.** Relative abundance of bacteria **(A)** phyla and **(B)** genera in the gut microbiota of each synbiotic treatment group. **(C)** Representative gel electrophoresis results of a polymerase chain reaction (PCR) analysis for the bile salt hydrolase (*bsh)* gene present in the *L. fermentum* K73 genome. Two time points were processed per treatment: day 0 (D0) before administration and day 21 (D21) after the start of treatment with synbiotic.

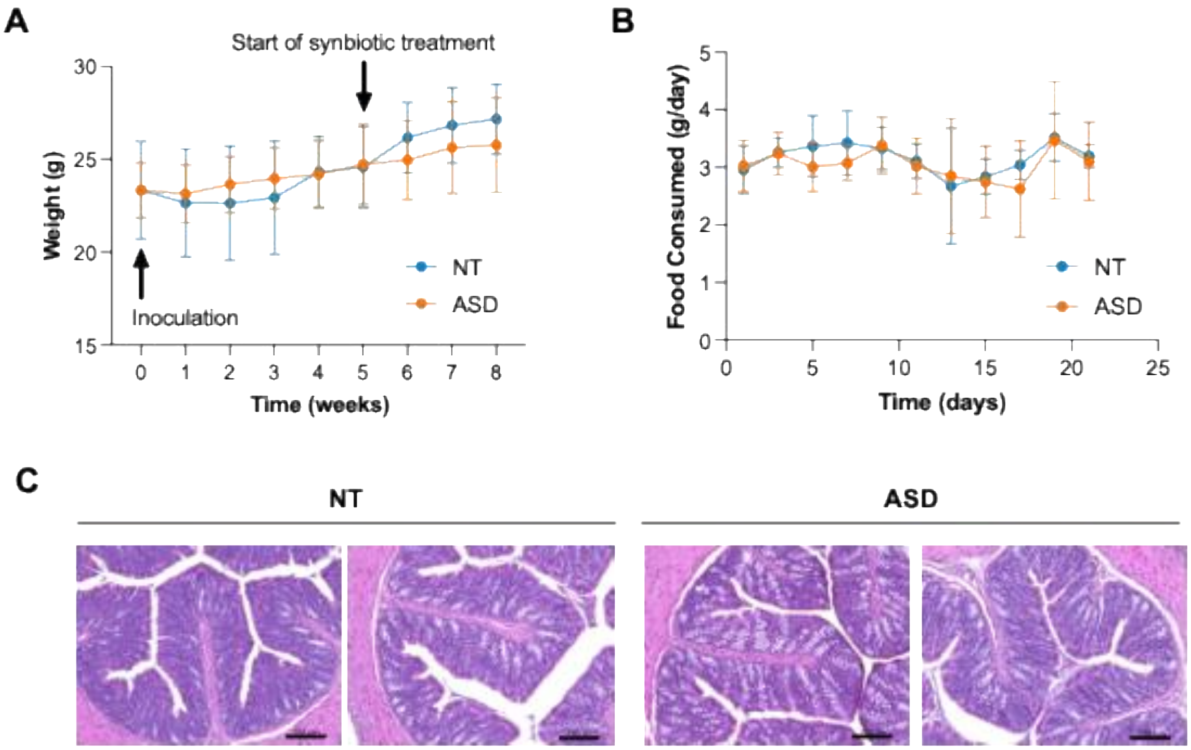


**Supplementary Figure 4. Synbiotic treatment did not have adverse health effects in GF mice transplanted with microbiota from NT and ASD donors. (A)** Average mouse body weight recorded weekly starting at inoculation with microbiota from NT or ASD donors. Data are presented as mean ± SD. **(B)** Average daily food consumption after the start of dietary supplementation with the synbiotic. Data are presented as mean ± standard deviation. **(C)** Representative images of H&E-stained colon sections obtained from two mice transplanted with fecal microbiota from two distinct donors per group. The colons were dissected 21 days after the start of synbiotic treatment. Scale bar represents 100 μm.

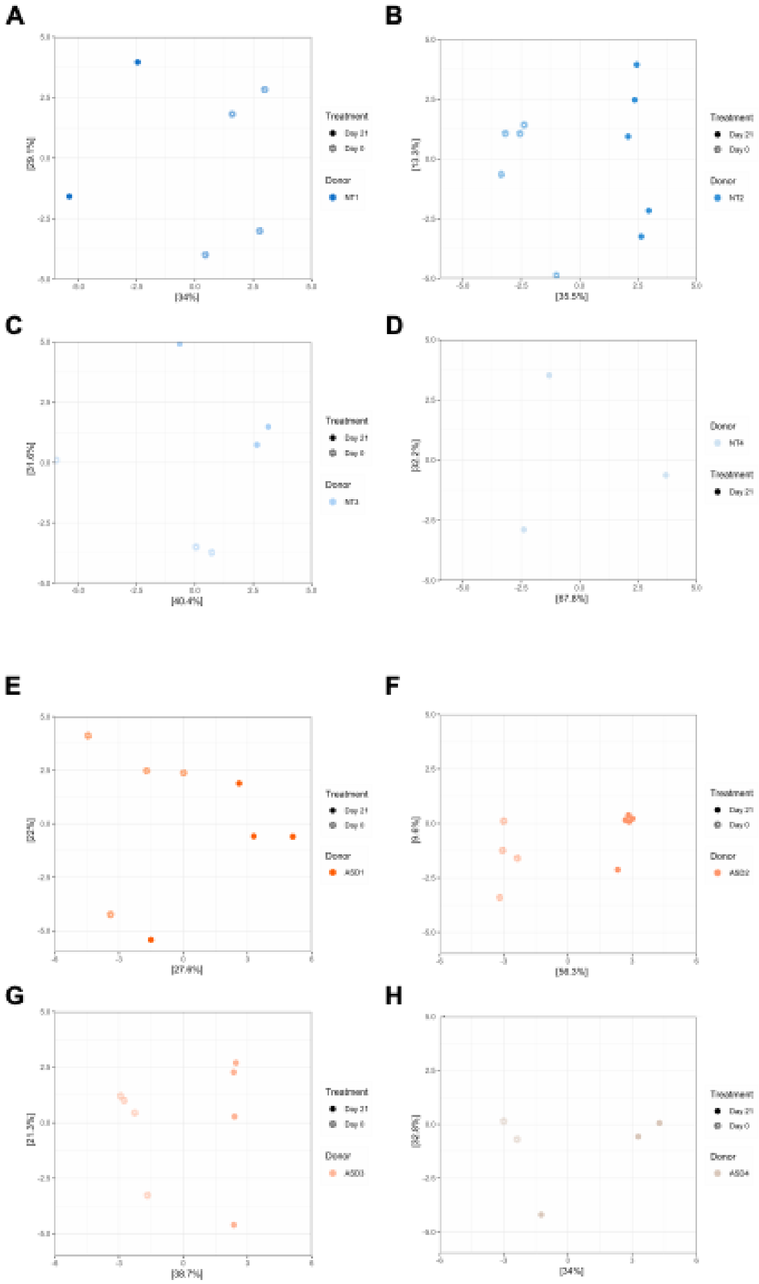


**Supplementary Figure 5. Analysis of beta diversity before and after synbiotic treatment by microbiota donor.** Individual PCA plots of Bray Curtis distances before (Day 0, open symbols) and after (Day 21, closed symbols) synbiotic treatment separated by the original microbiota (A-D) NT and (E-H) ASD donors.

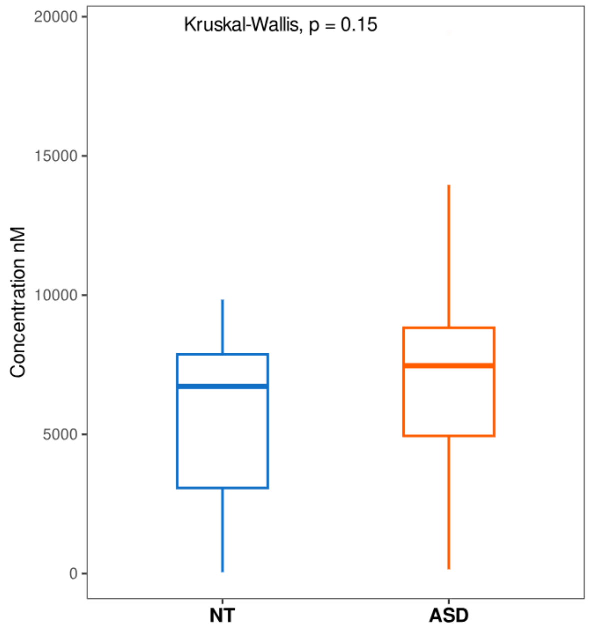


**Supplementary Figure 6. Quantification of serum serotonin levels via ELISA in mice colonized with NT or ASD derived microbiota.** Trunk blood was collected after behavioral analysis. Serum was processed according to manufacturer's guidelines, and no significant differences were observed. Statistical analysis was conducted using Kruskal-Wallis test ($p>0.05$).
